# Identification of key virus–prokaryote infection pairs that contribute to viral shunt in a freshwater lake

**DOI:** 10.1101/2023.02.05.527221

**Authors:** Shang Shen, Kento Tominaga, Kenji Tsuchiya, Tomonari Matsuda, Takashi Yoshida, Yoshihisa Shimizu

## Abstract

Viruses infect and kill productive prokaryotes in a density-or frequency-dependent manner and affect carbon cycling. However, the effects of the stratification transition, including the stratified and destratified periods, on the changes in prokaryotic/viral communities and the interactions among them remain unclear. We conducted a monthly survey of the surface and deep layers of a large and deep freshwater lake (Lake Biwa, Japan) for a year and analyzed the prokaryotic production and prokaryotic/viral metagenome. Our analysis (including 1 608 prokaryotes and 13 761 viruses) revealed that 19 prokaryotic species, accounting for ∼40% of total abundance, might be suppressed by viruses when prokaryotic production is higher. This suggests that a small proportion of prokaryotes contribute to a large amount of prokaryotic abundance, and these prokaryotes are infected and lysed by viruses, driving the viral shunt in the freshwater lake. Furthermore, we found that annual vertical mixing might yield a similar rate of community change between the surface and deep layers. This finding might be valuable in understanding how the communities change when the stratification of freshwater lakes is affected by global warming in the future.

## Introduction

Viral infections are critical factors in prokaryote mortality, suppressing the dominance of prokaryotic species and resupplying carbon and nutrients from prokaryotic cells to dissolved organic matter pools (viral shunts) [1, 2]. Recently, molecular biology techniques have been developed to reveal the diversity of prokaryotes and viruses [3, 4], the way viruses shape the prokaryotic community [5, 6] and affect carbon cycling [7] in various aquatic environments, including oceans and freshwater lakes. For example, prokaryotic communities in the deep layers of oceans, where prokaryotic production is less than that in the surface layer, are vastly different from those in the surface layer [8]. These differences result from environmental differences (e.g., sunlight, water temperature, and concentration of organic matter/nutrients). Previous studies with 10 years of ocean surveys indicated that the prokaryotic community in the surface layer (5 m) changes faster than that in the deep layer (150 m) [9, 10]. Unlike that in oceans, the water column in freshwater lakes is completely mixed vertically once or twice a year. This event causes changes in the prokaryotic/viral communities during the shift from the stratified to the de-stratified period. Previous studies have focused on the difference in prokaryotic/viral communities between the surface and deep layer or between the seasons in freshwater lakes [4, 11, 12]. However, no study has conducted a monthly survey targeting the surface and deep layer over a year and revealed how the prokaryotic/viral communities or their interactions in the surface and deep layer change with changes in the stratification of the water column.

The flux of viral shunt depends on the season and the depth of the water column because viruses infect prokaryotes more frequently when prokaryotic production is high [13–15]. This results from the density/frequency-dependent lytic infection cycle occurring between viruses and prokaryotes [16, 17]. The infection pairs between prokaryotes and viruses that contribute to the viral shunt are not known. Therefore, the prokaryotic species contributing to prokaryotic production should be investigated. Furthermore, whether these prokaryotic species are suppressed by their viruses should be determined. A previous study showed that monthly sampling can reveal a tight coupling of viruses and host prokaryotes [18]. However, it is still not known whether the host prokaryotes suppressed by viruses are related to prokaryotic production. Therefore, we considered that monthly sampling combined with prokaryotic production analysis can determine the infection pairs contributing to the viral shunt.

Lake Biwa, the largest lake in Japan, is a monomictic freshwater lake (surface area: 674 km^2^; maximum depth: 104 m). In the summer, higher prokaryotic production leads to higher viral infection in the surface layer, and approximately 70% of the production is reconnected to the dissolved organic matter pool via a viral shunt [19, 20]. During this season, the prokaryotic/viral communities in the surface and deep layers are entirely different [4, 21]. In the present study, we analyzed seasonal variation in prokaryotic production and prokaryotic/viral communities in both the surface and deep layers over the course of a year. Our goal was to clarify how stratification transition affects the communities and prokaryotic production in the surface/deep layers and which species contribute to the viral shunt.

## Materials and Methods

### Sample and data collection

Thirty-two lake water samples were collected at the pelagic station in Lake Biwa (35°23’
s41”N, 136°07’57”E, total depth of the sampling station: 90 m) between September 2018 and December 2019 (Table S1), from two depth layers: surface layer (depth: 0.5 m) and deep layer (depth: 60 m).

A dataset of environmental parameters (water temperature and dissolved oxygen (DO) concentration) was obtained from white papers published previously [22, 23].

### Analyses of prokaryotic production and prokaryotic/viral abundance

Prokaryotic production was measured using the [^15^N5]-2-deoxyadenosine (^15^N-dA)-incorporation method described earlier [19, 24]. The incorporation rate of ^15^N-dA (pmol L^−1^ d^−1^) was converted to the bacterial growth rate (cell L^−1^ d^−1^) using a conversion factor (1.83 × 10^6^ cells (pmol ^15^N-dA) ^−1^) determined in the same lake [19].

The collected lake water sample was fixed with buffered glutaraldehyde (pH 7.2, final concentration: 1%) immediately onboard, incubated at 4 °C before transportation to the laboratory, and stored at −30 °C. Slide glasses were prepared using DNA staining methods, [25] except that the SYBR Gold kit (Thermo Fisher Scientific, Oregon, USA; stock solution diluted 1:400) was used instead of the SYBR Green I kits [26]. Prokaryotic cells and virus-like particles were counted under an epifluorescence microscope (BZ-9000, KEYENCE) at 1 000 × magnification. Ten fields with at least 200 cells or virus-like particles were counted per sample. The slides were stored at −30 °C until counting.

### Partial 16S rRNA gene amplicon sequencing

The collected samples (250 mL) were prefiltered through GF/C filters (pore size: 1.2 μm, Merck). The filtrates were filtered through polycarbonate filters (pore size: 0.2 μm). DNA was extracted using a DNeasy PowerWater Kit (Qiagen, Hilden, Germany). The V3-V4 region of the 16S rRNA gene was amplified using the primer pair 341F (CCTACGGGNGGCWGCAG) and 805R (GACTACHVGGGTATCTAATCC) [27]. A sequencing library was prepared using the Illumina standard protocol (15044223 B, 16S Metagenomic Sequencing Library Preparation), and sequenced using a MiSeq sequencing system with a V3 reagent kit (300 × 2 bp, Illumina, San Diego, CA, USA).

The sequenced reads were demultiplexed and trimmed using Claident (v0.2.2019.05.10) by removing reads that included low-quality regions with default settings [28]. To differentiate amplicon sequence variants (ASVs), the trimmed reads were analyzed using the DADA2 pipeline (v. 1.16) in R version 3.5.1, with the settings of “trimLeft = 17, trimRight = 21” (Table S2) [29, 30]. ASVs were first classified using the ARB database [31] [32]. The unclassified ASVs were classified using Claident with default settings against the NCBI nucleotide sequence database [28, 33]. ASVs not assigned as prokaryotes (eukaryotes, mitochondria, and chloroplasts) were removed.

### Viral DNA preparation and sequencing

Four liters of collected lake water were prefiltered through GF/C filters (pore size: 1.2 μm) and then filtered through polycarbonate filters (pore size: 0.2 μm). The viral particles in the filtrates were concentrated using the Fe-based flocculation method and then purified using CsCl density centrifugation [34, 35]. Free DNA was removed via DNase treatment (Recombinant DNase I; Takara, Japan). Viral DNA was extracted using the DNeasy Blood & Tissue Kits (Qiagen, Hilden, Germany). A sequencing library was prepared according to the Illumina standard protocol (15031942 Rev. C, Nextera XT DNA Sample Preparation Guide), except that we used 0.25 ng viral DNA as an initial input [36]. The prepared library was sequenced using a MiSeq sequencing system with a V3 reagent kit (300 × 2 bp; Illumina, 4–6 samples). Lake Biwa viral contigs/genomes (LBVs) were constructed from the raw reads (see “Genome assembly, gene prediction, and an annotation” in supplementary information). Open reading frames were predicted using Prodigal v. 2.6.3 with the “-p meta” option [37]. The gene function of the predicted open reading frames was annotated according to a protocol mentioned in a previous study (see “Workflow for gene functional annotation” in the supplementary information [4]). The hosts of these viruses were predicted using multiple methods [38], as described in the section “Host prediction” in the supplementary information.

### Community data analysis

The relative abundance of LBVs was calculated as fragments per kilobase of contig per million mapped reads (FPKM) using the CountMappedReads2 script (https://github.com/yosuken/CountMappedReads2). The coverage of each LBV in each sample was calculated using CoverM (https://github.com/wwood/CoverM). The FPKM was set to zero when the coverage of the LBV was less than 80% of the genome/contig length [39]. The habitat preference of each LBV in each season is calculated as follows [4]:

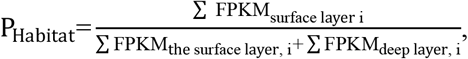

where i indicates months in certain seasons (e.g., i = 7, 8, and 9 in the middle of the stratified period, Table 1). P_Habitat_ of a LBV = 0 indicates that the LBV is exist only in the deep layer.

**Table 1.**
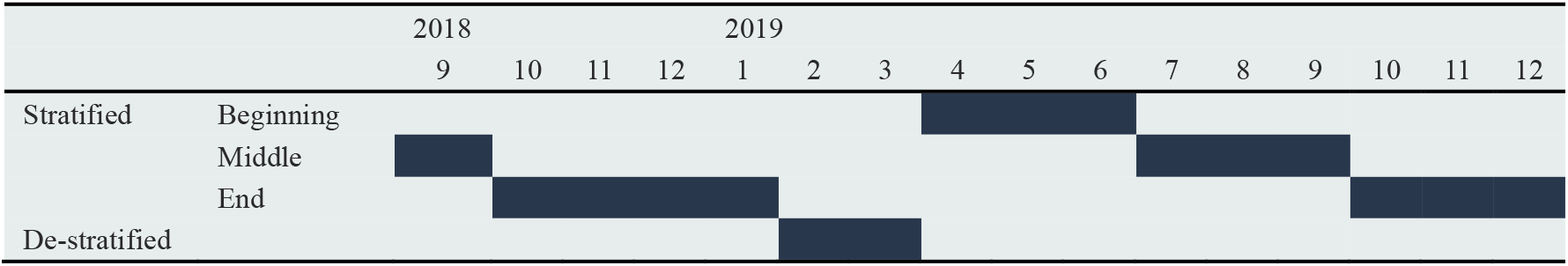
Basic information on season classification

Before community analysis, the dataset of each sample was rarefied based on coverage. The rarefied datasets were then used to calculate the Shannon index (*H’*) and generate non-metric multidimensional scaling (NMDS) plots. The dissimilarity of sample pairs (surface vs. deep layer, two samples with monthly separation) was calculated based on the Bray–Curtis dissimilarity. These community analyses were performed using the vegan v.2.5-7 package (https://CRAN.R-project.org/package=vegan) in R language [29].

For the dominance patterns of prokaryotic species (ASVs), the following three categories were defined: the persistent patterns were defined as ASVs dominated during >75% of the study period, the short pattern was defined as ASVs dominated during <10% of the study period (but at least 1 month) as described earlier [40], and the intermediate patterns were defined between the short and persistent patterns.

### Extraction of virus–prokaryote pairs related to viral shunt in the surface layer

To extract virus–prokaryote infection pairs that could contribute to the viral shunt in the surface layer during the summer (middle of the stratified period), when bacterial production is high (July–September 2019, Fig. 1F), we performed co-occurrence network analysis with ASV_dominant_ and LBVs. The LBVs those hosts were successfully predicted (phylum/class level) were applied to the co-occurrence network analysis. The ASV_dominant_ was defined as the trend of relative abundance that was the highest in the middle of the stratified period in 2019 and when the highest relative abundance was more than 1%. Infectious pairs between ASV_dominant_ and LBVs were successfully predicted and extracted using extended local similarity analysis [41, 42] via Pearson’s correlation test (*p* < 0.05, Q < 0.05).

**Fig. 1.**
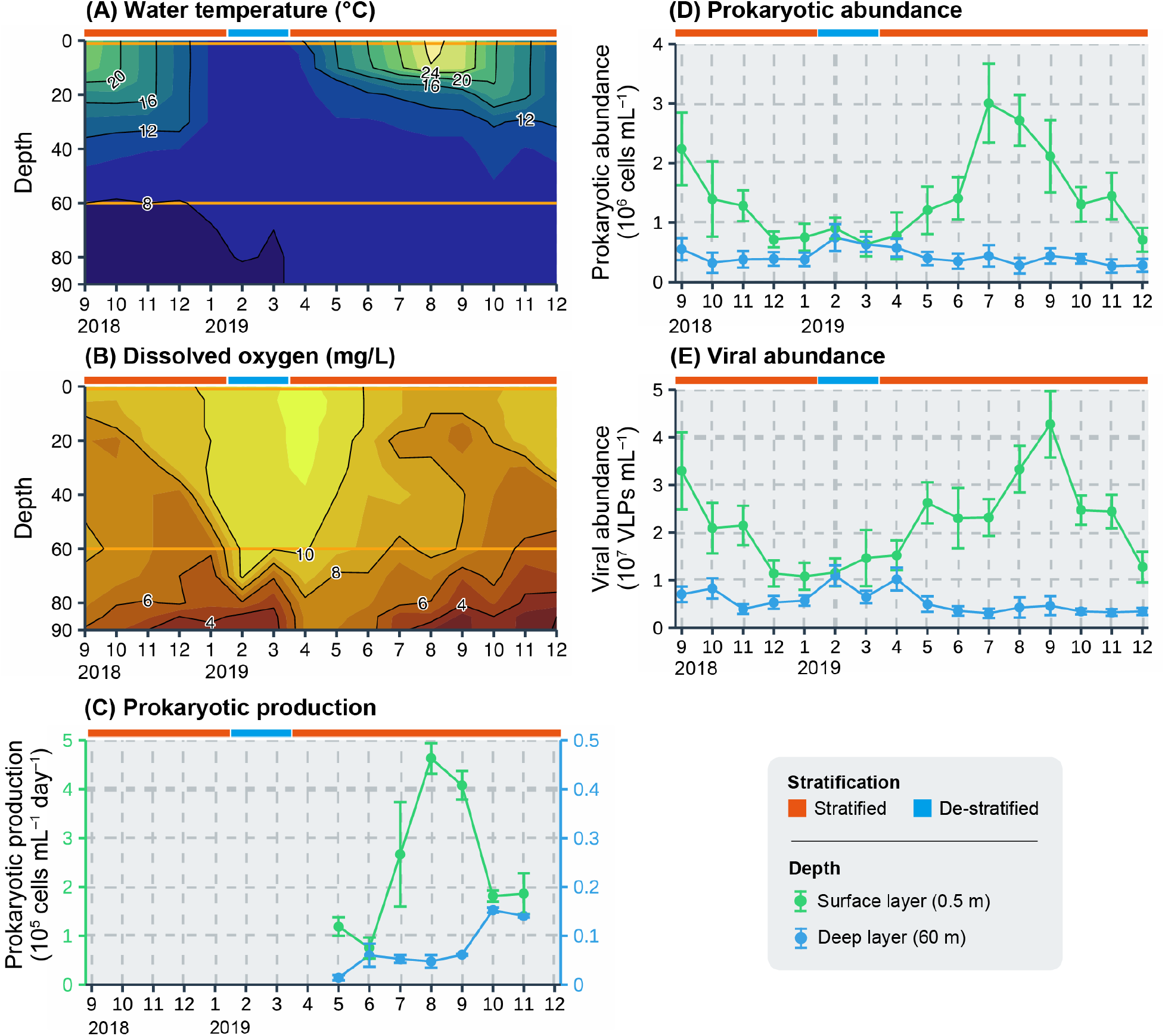
Seasonal variation in the environmental parameters through the study period:. (A) water temperature, (B) concentration of dissolved oxygen, (C) prokaryotic production, (D) prokaryotic abundance, and (E) viral abundance. Orange lines in panels (A) and (B) indicate the sampling depths (0.5 and 60 m). Values in panels (C), (D), and (E) represent the mean ± standard deviation (n = 3 in panel [C] and n = 10 in panel [D, E]).

### Data processing and visualization

Data handling was performed using the tidyverse v.1.3.1 package [43], and all figures were visualized using the ggplot2 v. 3.3.5 package [44] in R language. For the samples that met normality and equal variance assumptions, Student’s *t*-test was used to test the statistical significance of observed differences in the dissimilarity of time series separation. For the samples that did not meet these assumptions, the Mann–Whitney *U* test was applied. Results with *p* values < 0.05 were considered statistically significant.

## Results

### Water temperature and dissolved oxygen concentration

Water temperature in the surface layer decreased from 24.1 °C in September 2018 to 8.5 °C in February 2019 and increased again to 28.9 °C in August 2019, which was the highest in this study period (Fig 1A). In the deep layer, water temperature varied between 7.8 °C and 9.4 °C, which was lower and less variable than that in the surface layer. The de-stratified period, during which the water column was vertically and completely mixed, was defined as the period when the water temperature was the same in the surface and deep layer (<1 °C), and the DO was supplied to the deep layer. In the present study, we considered that February and March represented the destratified period (10.4–10.9 mgO_2_/L in the surface layer and 10.1– 10.3 mgO_2_/L in the deep layer, Fig 1B), whereas the other months represented the stratified period. Moreover, we divided the stratified period into the following three parts (Table 1): the beginning of the stratified period (April to June 2019), the middle of the stratified period (September 2018 and July to September 2019), and the end of the stratified period (October 2018 to January 2019 and October to December 2019). Notably, during this study period, incomplete vertical mixing was first observed during winter (January to March) since regular monitoring began in Lake Biwa in 1979 [45]. However, in February 2019, DO was supplied to a depth of 60 m, which was the sampling depth in this study.

### Prokaryotic/viral abundance and prokaryotic production

Prokaryotic production in the surface layer increased with the water temperature and reached 4.6 × 10^5^ cells mL^−1^ d^−1^ in the middle of the stratified period (August 2019, Fig. 1C). Prokaryotic production in the deep layer varied at 0.014–0.15 × 10^5^ cells mL^−1^ d^−1^ (median: 0.06 × 10^5^ cells mL^−1^ d^−1^) and was lower and less variable than that in the surface layer. Prokaryotic production in the surface layer was 12–97-fold higher than that in the deep layer.

Prokaryotic and viral abundances in the surface layer (prokaryotes: 0.64–3.0 × 10^6^ cells mL^−1^, viruses: 1.1–4.3 × 10^7^ VLPs mL^−1^) were the highest during the middle of the stratified period, when the water temperature is also higher than that during the other seasons (Fig. 1D and E). These three parameters were less variable and lower in the deep layer (prokaryotes: 0.27–0.75 × 10^6^ cells mL^−1^, viruses: 0.31– 1.1 × 10^7^ VLPs mL^−1^) than in the surface layer. The prokaryotic/viral abundance in the surface and deep layers showed similar values when the de-stratified period began (i.e., February).

### Diversity and seasonal variation of prokaryotic communities

A total of 1 608 ASVs were detected from 1 497 727 non-chimeric reads obtained from 32 samples using 16S rRNA gene amplicon sequencing (Table S2). Actinobacteria was the most predominant phylum in both the surface and deep layers throughout the study period (Fig. S1A). Bacteroidetes and Cyanobacteria were also predominant in the surface layer, whereas Chloroflexi and Verrucomicrobia were predominant in the deep layer. According to the Bray–Curtis dissimilarity comparison between the surface and deep layers, the two communities were most closely related during the de-stratified period (0.084 in February 2019) and most different during the middle of stratified period (0.90 in September 2019, Fig. 2A). The NMDS analysis showed that the community in the surface layer was more variable than that in the deep layer (Fig. S2A).

**Fig. 2.**
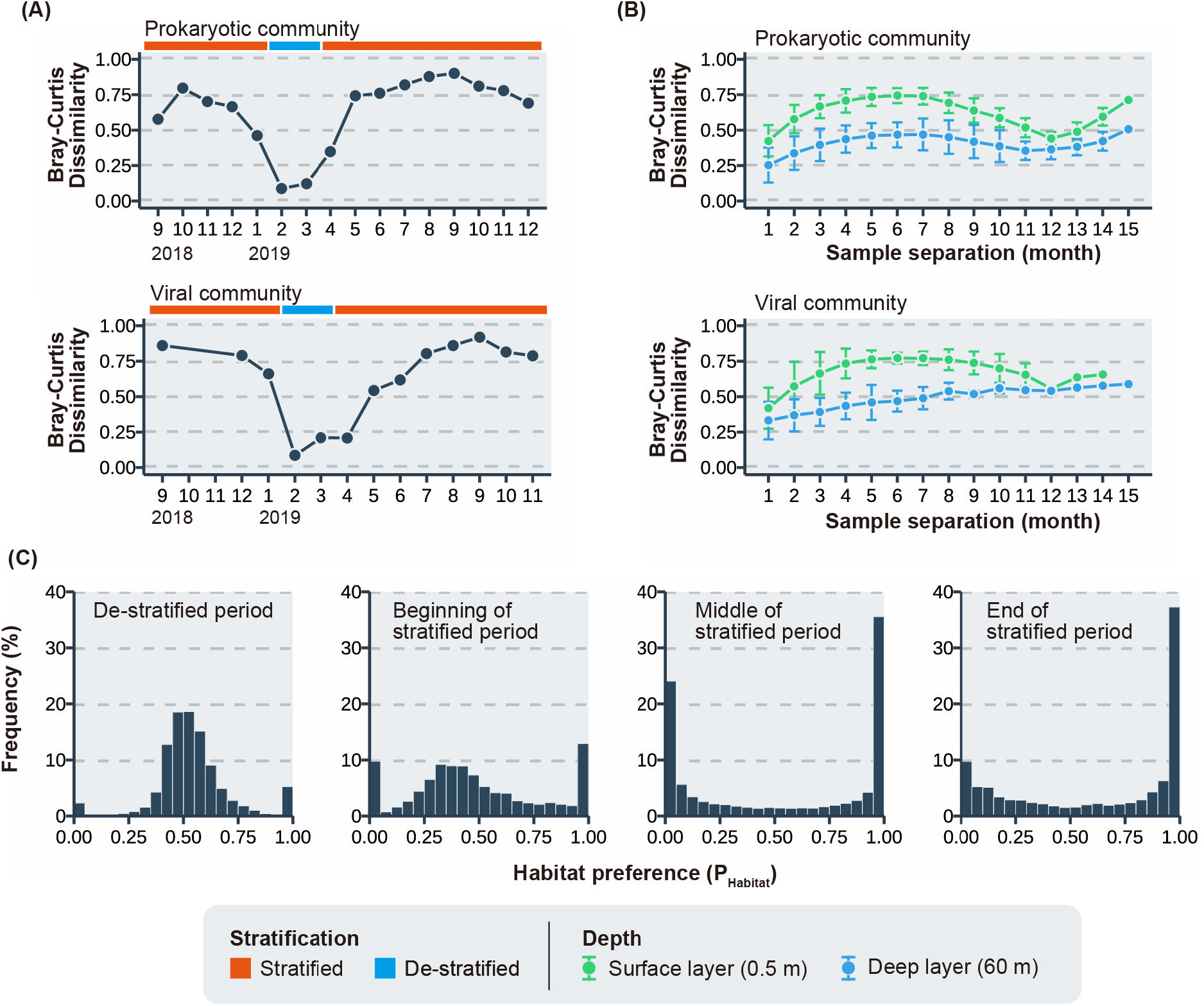
Community analysis for prokaryotes and viruses. (A) P_Habitat_ of LBVs. The sampling period was classified into four seasons, as described in Table 1. (B) LBV of P_Habitat_ is 0, which means that LBV exists only in the deep layer. (B) Bray–Curtis dissimilarity of prokaryotic/viral community between the surface and deep layers. (C) Bray–Curtis dissimilarity of sample separation (month) for prokaryotic/viral communities. Each plot and error bar reflects the mean ± standard deviation. The results of the statistical analysis are shown in Table S5. P_Habitat_ indicates habitat preference, and LBVs indicate Lake Biwa viruses.

According to the dissimilarity comparing two samples based on the time (month) of separation, the dissimilarity in the two layers showed a local maximum around 6 months later (0.75 and 0.48 in the surface and deep layer, respectively) and a local minimum 12 months later (0.47 and 0.40 in the surface and deep layer, respectively; Fig. 2B). The dissimilarities of the prokaryotic communities in the surface layer after 6 months were significantly higher than those in the deep layer, and the dissimilarities after 12 months in the two layers showed no significant difference (Table S5).

### Diversity and seasonal variation of viral communities

A total of 13 761 LBVs were detected in 134 496 816 high-quality paired-end reads from the 27 samples (Table S3). The 385 LBVs had complete (i.e., circular) genomes. A total of 2 780 LBVs successfully predicted their hosts at the phylum/class level (Table S4, 10.6–29.5% of total LBVs FPKM-based abundance). Of these LBVs, viruses specific to members from the phyla Actinobacteria and Cyanobacteria were predominant in the surface layer, whereas those specific to members from the phyla Actinobacteria, Gammaproteobacteria, and Verrucomicrobia were predominant in the deep layer (Fig. S1B). The diversity of the viral community (Shannon *H*’) was higher than that of the prokaryotic community (Fig. S3).

P_Habitat_ in the de-stratified period showed a distribution with a center of 0.5, whereas the P_Habitat_ in the middle of the stratified period showed a distribution with both ends (< 0.05, > 0.95; Fig. 2C).

According to the Bray–Curtis dissimilarity, between the surface and deep layers, the dissimilarity was the highest in the middle of the stratified period (0.86 in September 2018/2019) and the lowest in the de-stratified period (February 2019: 0.085; Fig. 2A). The NMDS analysis showed that the community in the surface layer was more variable than that in the deep layer (Fig. S2B). According to the dissimilarity comparing two samples based on the time (month) of separation, the dissimilarity in the surface layer showed a local maximum 6 months later (0.77) and a local minimum 12 months later (0.56), whereas in the deep layer, the dissimilarity monotonically increased throughout the study period. The dissimilarities of the viral communities in the surface layer after 6 months were significantly higher than those in the deep layer (Table S5). After 12 months, the dissimilarity showed similar values in the surface and deep layer (0.56 and 0.54, respectively).

### Virus–prokaryote infection pairs related to viral shunt in the surface layer

A total of 21 ASVs (ASV_dominant_, relative abundance is > 1% during the middle of the stratified period), including those for Actinobacteria, Alphaproteobacteria, Betaproteobacteria, Bacteroidetes, Cyanobacteria, and Verrucomicrobia, showed the highest relative abundance (>1%) when the prokaryotic production was high (Fig. 3, Table S6). The maximum relative abundance of these ASV_dominant_ varieties varied from 1.0 to 14.1%. The total relative abundance of these 21 ASV_dominant_ was 41.3% in July, 56.3% in August, and 46.2% in September. For 19 ASV_dominant_ of the 21 ASV_dominant_, at least one LBV co-occurred (Fig S4). Of the remaining 2 ASV_dominant_, LBVs co-occurring with ASV_41 (Phylum Chlorobi) could not be detected and ASV_204 (Phylum Gemmatimonadetes) was excluded from the analysis because its phages could not be predicted. In total, 396 LBVs were found to co-occur with these 19 ASVs, accounting for 4.3%, 10.8%, and 7.6% in July, August, and September, respectively, as determined based on the FPKM-based relative abundance (Table S6). Of the 19 ASV_dominant_, 13 ASV_dominant_ species were undetectable or showed low abundance before their abundance increased in the middle of the stratified period, and they consisted of Actinobacteria, Bacteroidetes, Cyanobacteria, Proteobacteria (alpha and beta), and Verrucomicrobia. The remaining 6 ASV_dominant_ (ASV_4, 5, 7, 80, 84, 122) were also dominant in other periods.

**Fig. 3.**
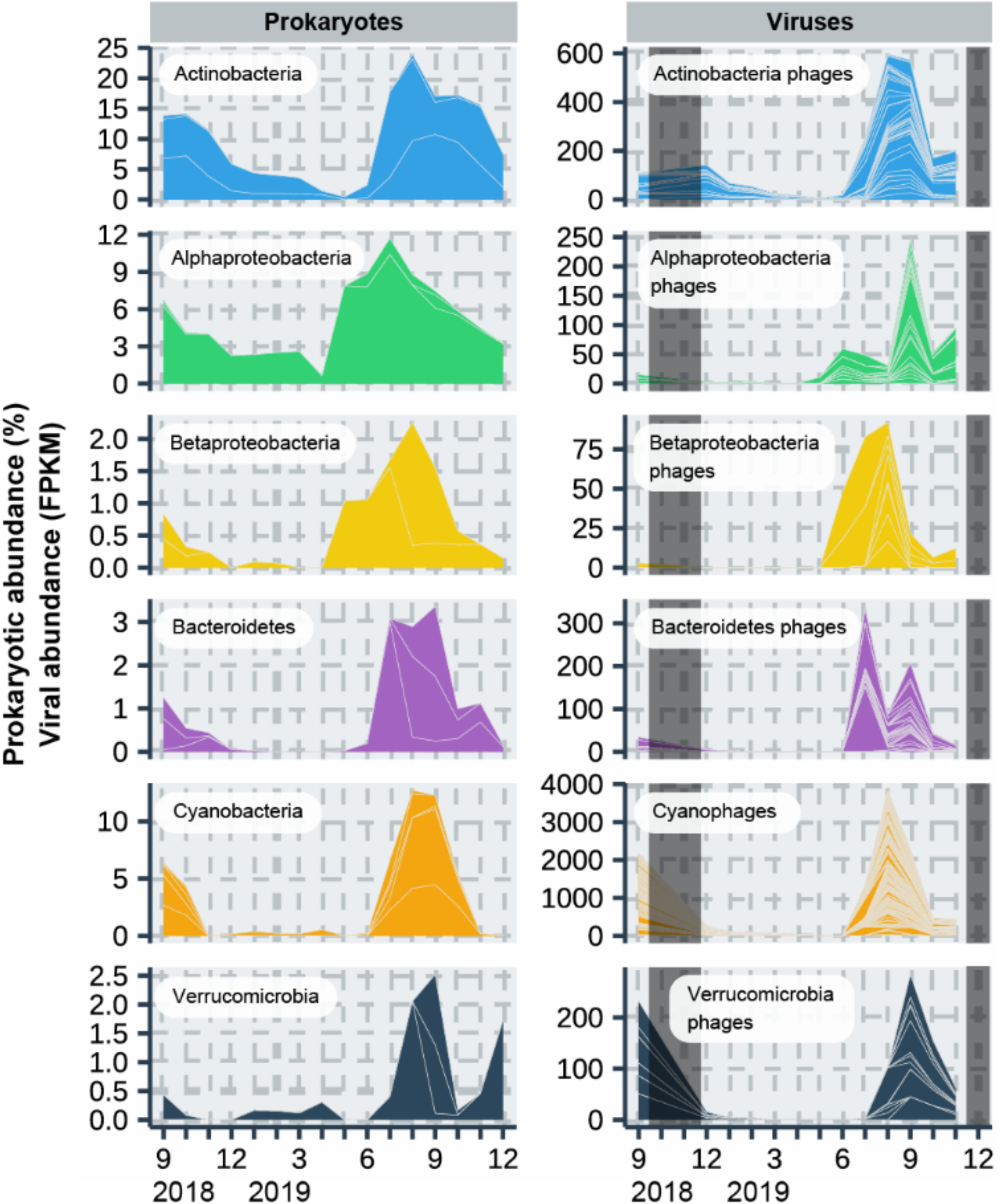
Seasonal variation in the abundance of prokaryotic species (19 ASVs) may contribute to prokaryotic production and LBVs co-occurring with the ASVs in the surface layer. Gray shading indicates no data (Table S1). Individual plots for ASVs and co-occurring LBVs are shown in Fig. S4. ASVs indicate amplicon sequence variants, and LBVs indicate Lake Biwa viruses.

### Dominant patterns of the prokaryotic/viral community

The number of ASVs with relative abundances of more than 1% for at least one month during this study period was 99 and 65 in the surface and deep layers, respectively (Fig. 4A), corresponding to approximately 5% of the total ASVs (i.e., 1 608 species) detected. Of these, in the surface layer, approximately 44 (40%) and 5 (5%) of the ASVs showed short (i.e., dominant for 1 month) and persistent (i.e., dominant for more than 12 months) patterns, respectively (Fig. 4A). Most of these existed (>0%) for less than 8 months (e.g., ASV_89, 169, 177, and 305 in Fig. S4). In the deep layer, approximately 23 (35%) and 15 (20%) of the ASVs showed short and persistent patterns, respectively. Furthermore, 50% of the dominant ASVs in the deep layer existed throughout the study period (16 months).

**Fig. 4.**
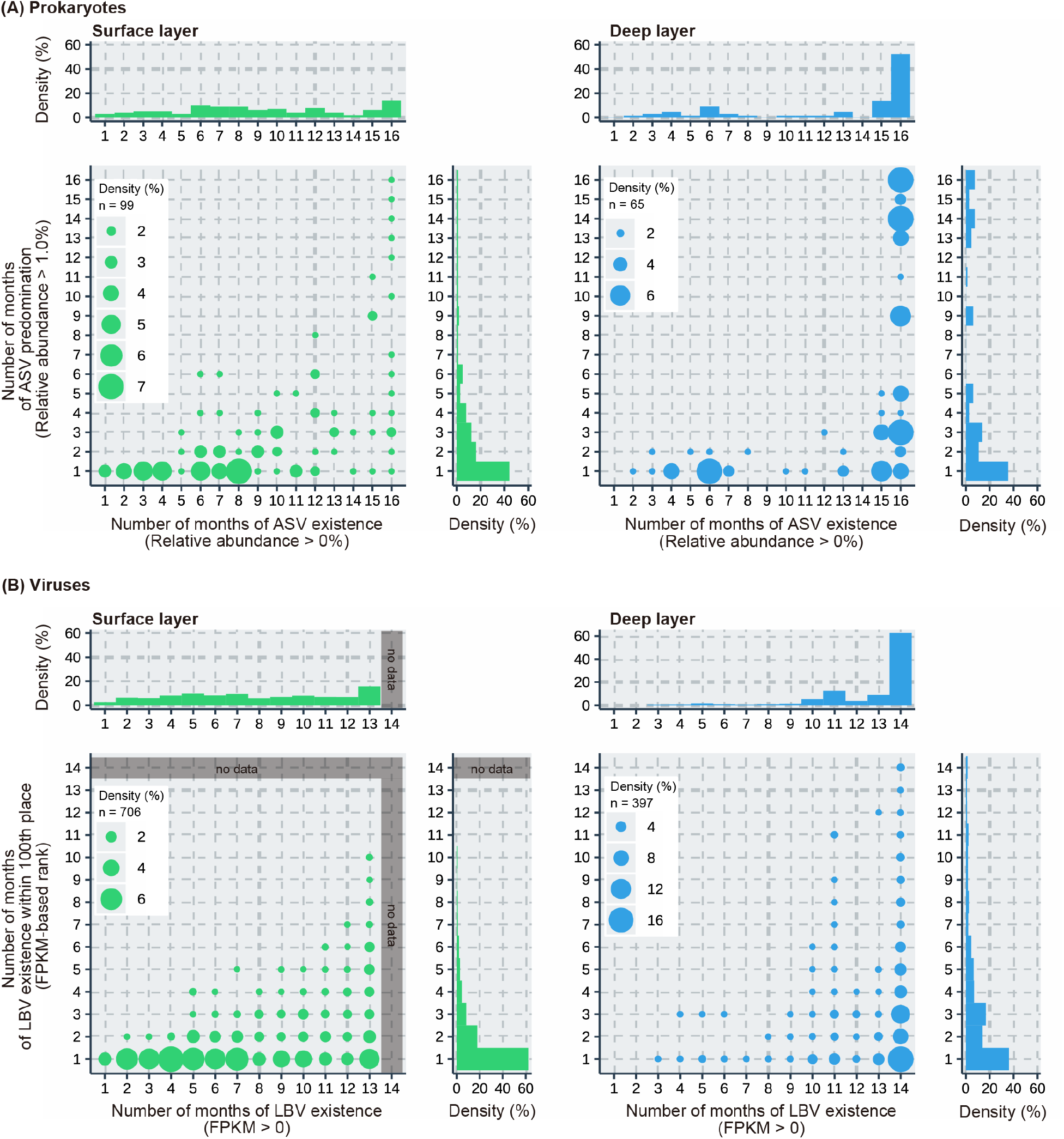
Existence vs. predominance plots for (A) prokaryotes and (B) viruses in the surface and deep layers. The X-axis indicates the number of months that amplicon sequence variant (ASV)/Lake Biwa virus (LBV) existed during the study period. The Y-axis indicates the number of months for which ASV/LBV predominated. Predominant ASV is defined as that whose relative abundance is higher than 1%. Predominated LBV is defined as that with an FPKM-based rank within 100 (13 761 LBVs were detected in this study). ASVs indicate amplicon sequence variants, and LBVs indicate Lake Biwa viruses.

Dominant LBVs, those ranked within 100 based on the FPKM data in the surface layer, showed a pattern similar to that of ASVs in the surface layer (Fig. 4B). Approximately 60% of LBVs showed a short pattern. In the deep layer, the dominant patterns of LBVs were different from those of ASVs. Approximately 40% of LBVs showed a short pattern but fewer LBVs showed a broad pattern. Moreover, 60% of dominant LBVs in the deep layer existed (FPKM >0) throughout the study period, but only 20% of dominant LBVs in the surface layer existed throughout the study period.

## Discussion

The data we obtained via annual sampling from the surface and deep layers of the freshwater lake led to two major findings. First, a small number of prokaryotes, accounting for approximately 50% of the abundance, may be infected and suppressed by viruses when prokaryotic production was high and contribute to the viral shunt.

Second, annual vertical mixing of the water column may result in a similar rate of community change between the surface and deep layers.

### The viral shunt in summer may be exclusively driven by a few prokaryotic species

In the middle of the stratified period (Summer), prokaryotic production and prokaryotic and viral abundance were the highest in the surface layer throughout the study period (Fig. 1). The viral infection rate is also the highest in this period, and approximately 40–60% of prokaryotes are reconnected to the pool of dissolved organic matter via viral lysis [20, 46]. One of the major successes of this study was that we estimated the potential virus–prokaryote infection pairs contributing to the viral shunt. Infection pairs, including 19 ASV_dominant_ and 396 LBVs, were successfully estimated in the middle of the stratified period. These 19 ASV_dominant_ accounted for 41.3–56.3% of total prokaryotic abundance. Of the 19 ASV_dominant_, 13 species were undetectable or showed low abundance in other periods but could grow more actively (*r*-strategy) than other ASVs in response to the increase in water temperature or dissolved organic matter derived from primary production at the middle of the stratified period. This is supported by the findings of previous studies [47–52]. For instance, the growth rate and absolute abundance of *Synechococcus* (GpIIa) increased with the increase in water temperature [50, 51]. Actinobacteria and proteobacteria increased in their relative abundances with the increase in the labile substrate [49], and Verrucomicrobia could rapidly respond to carbohydrates or cyanobacterial bloom [47, 48, 52]. Concentrations of these labile substances are higher in the surface layer during the middle of the stratified period in Lake Biwa [53]. Thus, these 13 ASV_dominant_ might be growing actively, thereby contributing to prokaryotic production, and can be infected by viruses based on the density- and frequency-dependent models [16, 17]. In this study, bacteria that do not incorporate dA (lack of deoxyribonucleoside kinases) were not included in the prokaryotic production analysis [54]. Moreover, minor (<1% of relative abundance) but active bacteria were not considered the ASV_dominant_ in this study. However, a previous study using the bromodeoxyuridine-incorporating method reported that approximately 60% of prokaryotic cells showed growing activity in oligotrophic Lake Stechlin (Germany) [55], which is consistent with our result that the 19 ASV_dominant_ accounted for 41.3–56.3% of total prokaryotic abundance. These results supported that a few prokaryotic species drive most part of the viral shunt during the middle of the stratified period. Our results do not deny the possibility of grazing pressure on these 19 ASVs in the middle of the stratified period. Protists also actively graze prokaryotes in the middle of the stratified period in Lake Biwa [56], and these 19 ASVs may have also been grazed by protists. In this study, we only measured the prokaryotic production of free-living prokaryotes in the GF/C filtrates but not the prokaryotic production or interaction between prokaryotes and viruses attached to the particulate organic matter.

Because viruses may affect the dynamics of particulate organic matter [57, 58], further studies should be conducted, such as determining which viral group may affect particle export.

In the deep layer, the number of ASVs with co-occurring viruses was lower than that in the surface layer (82 and 26 ASVs in the surface and deep layers, respectively). First, because the prokaryotic/viral community in the deep layer did not change dynamically, fewer infection pairs were detected using co-occurrence network analysis. Second, approximately 20% of LBVs could be predicted by their host phylum/class, and the remaining 80% remained unknown. LBVs in the deep layer (also in the surface layer) that were not detected to co-occur with ASVs could be included in the remaining LBVs. Third, oligotrophic environments (lower host abundance or prokaryotic production) favor lysogenic viruses, whereas lytic viruses are dominant in highly productive environments [59, 60]. In the deep layer, prokaryotic production was markedly lesser than that in the surface layer (Fig. 1C), and the infection strategy of some viruses in the deep layer might be lysogeny. Furthermore, viruses in the deep layer could be lysogenic when their host becomes abundant based on the Piggyback-the-Winner model [61]. However, of the 397 LBVs that ranked within 100 based on the FPKM data, only 14 LBVs carried the integrase gene (Table S9). To test this possibility, further analyses, such as analysis of the viral metagenome in prokaryotic cells, must be carried out [4].

### Most prokaryotes and viruses in the deep layer persist during all months but dominate in only one month

In the surface layer, environmental parameters, such as water temperature and the phytoplankton community composition [62], change dynamically, resulting in changes in the composition of organic matter. This kind of rapid environmental change may favor approximately 40% of dominant ASV species, which supports the observed short patterns of abundant ASVs (relative abundance >1%) in the surface layer. Viruses also showed a pattern similar to that of ASVs in the surface layer, reflecting that these viruses increased in abundance by infecting and lysing their host prokaryotes.

In the deep layer, both patterns were observed for the dominant ASVs and unlike the surface layer, more ASVs showed persistent patterns (i.e., >12 months). This can be explained by the environmental stability of deep layers. In the deep layer, water temperature is stable (7.8–9.4 °C), and prokaryotic production is notably lower than that in the surface layer (Fig. 1A, C). Moreover, new organic production is limited [63] and usable (labile) organic matter is depleted [20], leading to the slow growth of prokaryotes. Notably, 23 ASVs in the deep layer showed a short pattern (Fig. 4A). Of these, 7 ASVs increased during the de-stratified period and may have been supplied by the surface layer via vertical water mixing (e.g., ASV_11, 81, and 43 in Fig. S5). The other 16 ASVs (e.g., ASV_64 and 149 in Fig. S5) may increase in response to the substrate that may be provided by the surface layer through sinking particles or from lysates via infection of other ASVs. Interestingly, unlike the dominant ASVs, most of the dominant LBVs showed a short or intermediate (2–3 months) pattern, suggesting the result of ASVs with a short pattern. Moreover, most of these LBVs were present throughout the study period (i.e., 14 months). These results indicated that although the environment in the deep layer does not change considerably, some ASVs may occasionally become dominant and be suppressed by their phages, resulting in their phages ranking within 100. These dominant viruses in the deeper layer exist for a long time (10–14 months) probably because factors that remove viral particles—such as the particulate organic matter on which viruses attach or heterotrophic protists, which may ingest viral particles—are depleted. This could explain that how ASV composition changed seasonally but LBV composition changed monotonically in the deep layer (Fig. 2B). Another possibility is that some dominant ASVs (> 1%) are not infected by viruses. A previous study suggested that host species at an abundance of less than 10^4^ cells/mL are not infected by their phages [64]. The prokaryotic abundance in the deeper layer was less than 10^6^ cells/mL (Fig. 2D), and some dominant ASVs may not fulfill this effective infection threshold (i.e., 10^4^ cells/mL) even though their relative abundance was > 1%. This may result in the viral community in the deeper layer changing monotonically.

### The rate of community changes is differentiated by stratification and is uniform annually owing to vertical mixing

The prokaryotic/viral communities between the surface and deep layers were substantially different in the middle of the stratified period and mixed during the de-stratified period (Fig. 2A). This may have been affected by the differences in community changes between the two layers. The higher dissimilarities of the prokaryotic and viral communities in the surface layer 6 months later than those in the deep layer indicated that the communities in the surface layer changed faster than those in the deep layer. The 6-month time gap leads to opposite seasons being considered (e.g., spring vs. autumn, summer vs. winter), and these differences in dissimilarity may result from differences in prokaryotic production or environmental parameters.

For the prokaryotic/viral communities, the dissimilarity between the surface and deep layers was not substantially different after 12 months (Fig. 2B, Table S5). Interestingly, the gap in community changes between the surface and deep layers closed after 12 months, although prokaryotic production and environmental parameters between the two layers were different. These results indicated that prokaryotic and viral communities in the surface and deep layers change at a similar rate over the years. This finding is inconsistent with that of previous research wherein the prokaryotic community in the surface layer (5 m) was shown to change faster than that in the deep layer (150 m) of the ocean [9, 10]. Annual vertical mixing may explain why, unlike the ocean, community changes between the surface and deep layers were similar over years in Lake Biwa. Vertical mixing occurs each year (mostly from January to March) in Lake Biwa. In the de-stratified period, not only the abundance and composition of microorganisms but also environmental parameters (e.g., concentration of nutrients or organic matter) are vertically mixed and become similar throughout the water column ([22, 23]). Although rapid changes in prokaryotic production or environmental parameters in the surface layer result in faster changes in the prokaryotic/viral community, community changes are reset by vertical mixing in the de-stratified period every year. This could lead to a similar rate of change in prokaryotic/viral communities between the surface and deep layers after 12 months. Unlike in oceanic environments [9, 10], prokaryotic/viral communities in the deep layers of aquatic environments with vertical mixing may change at a rate similar to that in the surface layer.

Our monthly survey combining the community analysis and prokaryotic production revealed the way through which dominant prokaryotes are suppressed by a viral infection and identified the key infection pairs contributing to the viral shunt in the freshwater lake. The results of this study will help to understand how viruses affect carbon cycling via viral infections in prokaryotes that respond to primary production. Our analysis also indicated that the rate of community changes is similar between the surface and deep layers due to the annual vertical mixing even though the environments or prokaryotic activity in the surface/deep layer are different. This finding might aid in understanding the effect of climate change on stratification and the microbial community in the future.

## Acknowledgments

We thank the Lake Biwa Environmental Research Institute for assistance with field sampling. We also thank SuperComputer System, Institute for Chemical Research, Kyoto University for providing computer time. This research was supported by JSPS KAKENHI (grant numbers JP19J14985, JP20H04323, JP20K12140, JP21H05057, JP22K14351 and JP22J01607), Kurita Water and Environment Foundation (19B045), and Kyoto University Foundation.

## Data availability statement

Raw reads of the 16S rRNA genes from prokaryotic fractions have been deposited in GenBank (SAMD00576609-SAMD00576640). Raw reads of the viral fractions have been deposited in GenBank (SAMD00576528-SAMD00576554).

## References

1. Suttle CA. Viruses in the sea. Nature 2005; 437: 356‒361.

2. Suttle CA. Marine viruses - Major players in the global ecosystem. Nat Rev Microbiol 2007; 5: 801‒ 812.

3. Brum JR, Ignacio-Espinoza JC, Roux S, Doulcier G, Acinas SG, Alberti A, et al. Patterns and ecological drivers of ocean viral communities. Science (1979) 2015; 348: 1261498.

4. Okazaki Y, Nishimura Y, Yoshida T, Ogata H, Nakano S. Genome-resolved viral and cellular metagenomes revealed potential key virus-host interactions in a deep freshwater lake. Environ Microbiol 2019; 21: 4740‒4754.

5. Yoshida T, Nishimura Y, Watai H, Haruki N, Morimoto D, Kaneko H, et al. Locality and diel cycling of viral production revealed by a 24 h time course cross-omics analysis in a coastal region of Japan. ISME J 2018; 12: 1287‒1295.

6. Carlson MichaelCG, Ribalet F, Maidanik I, Durham BP, Hulata Y, Ferrón S, et al. Viruses affect picocyanobacterial abundance and biogeography in the North Pacific Ocean. Nat Microbiol 2022; 7: 570‒ 580.

7. Guidi L, Chaffron S, Bittner L, Eveillard D, Larhlimi A, Roux S, et al. Plankton networks driving carbon export in the oligotrophic ocean. Nature 2016; 532: 465‒470.

8. Hurwitz BL, Brum JR, Sullivan MB. Depth-stratified functional and taxonomic niche specialization in the ‘core’ and ‘flexible’ Pacific Ocean Virome. ISME J 2015; 9: 472‒484.

9. Fuhrman JA, Cram JA, Needham DM. Marine microbial community dynamics and their ecological interpretation. Nat Rev Microbiol 2015; 13: 133‒146.

10. Chow C-ET, Sachdeva R, Cram JA, Steele JA, Needham DM, Patel A, et al. Temporal variability and coherence of euphotic zone bacterial communities over a decade in the Southern California Bight. ISME J 2013; 7: 2259‒2273.

11. Moon K, Kim S, Kang I, Cho J-C. Viral metagenomes of Lake Soyang, the largest freshwater lake in South Korea. Sci Data 2020; 7: 349.

12. Coutinho FH, Cabello-Yeves PJ, Gonzalez-Serrano R, Rosselli R, López-Pérez M, Zemskaya TI, et al. New viral biogeochemical roles revealed through metagenomic analysis of Lake Baikal. Microbiome 2020; 8: 163.

13. Steward GF, Smith DC, Azam F. Abundance and production of bacteria and viruses in the Bering and Chukchi Seas. Mar Ecol Prog Ser 1996; 131: 287‒300.

14. Pradeep Ram AS, Mari X, Brune J, Torréton JP, Chu VT, Raimbault P, et al. Bacterial-viral interactions in the sea surface microlayer of a black carbon-dominated tropical coastal ecosystem (Halong Bay, Vietnam). Elementa: Science of the Anthropocene 2018; 6.

15. Weinbauer MG, Höfle MG. Significance of viral lysis and flagellate grazing as factors controlling bacterioplankton production in a eutrophic lake. Appl Environ Microbiol 1998; 64: 431‒438.

16. Thingstad TF. Elements of a theory for the mechanisms controlling abundance, diversity, and biogeochemical role of lytic bacterial viruses in aquatic systems. Limnol Oceanogr 2000; 45: 1320‒ 1328.

17. Rodriguez-Brito B, Li L, Wegley L, Furlan M, Angly F, Breitbart M, et al. Viral and microbial community dynamics in four aquatic environments. ISME J 2010; 4: 739.

18. Tominaga K, Ogawa-Haruki N, Nishimura Y, Watai H, Yamamoto K, Ogata H, et al. Prevalence of viral frequency-dependent infection in coastal marine prokaryotes revealed using monthly time series virome analysis. bioRxiv 2021; 2021.09.23.461490.

19. Tsuchiya K, Tomioka N, Sano T, Kohzu A, Komatsu K, Imai A, et al. Decrease in bacterial production over the past three decades in the north basin of Lake Biwa, Japan. Limnology (Tokyo) 2020; 21: 87‒96.

20. Shen S, Shimizu Y. Seasonal Variation in Viral Infection Rates and Cell Sizes of Infected Prokaryotes in a Large and Deep Freshwater Lake (Lake Biwa, Japan). Front Microbiol. 2021., 12: 1004

21. Okazaki Y, Nakano S. Vertical partitioning of freshwater bacterioplankton community in a deep mesotrophic lake with a fully oxygenated hypolimnion (Lake Biwa, Japan). Environ Microbiol Rep 2016; 8: 780‒788.

22. Shiga-Prefecture. Environmental white paper in 2019 (reference). 2019; 1‒202.

23. Shiga-Prefecture. Environmental white paper in 2020 (reference). 2020; 1‒205.

24. Tsuchiya K, Sano T, Kawasaki N, Fukuda H, Tomioka N, Hamasaki K, et al. New radioisotope-free method for measuring bacterial production using [15N5]-2’-deoxyadenosine and liquid chromatography mass spectrometry (LC‒MS) in aquatic environments. J Oceanogr 2015; 71: 675‒683.

25. Patel A, Noble RT, Steele JA, Schwalbach MS, Hewson I, Fuhrman JA. Virus and prokaryote enumeration from planktonic aquatic environments by epifluorescence microscopy with SYBR Green I. Nat Protoc 2007; 2: 269‒276.

26. Shibata A, Yoichi G, Saito H, Kikuchi T, Toda T, Taguchi S. Comparison of SYBR Green I and SYBR Gold Stains for Enumerating Bacteria and Viruses by Epifluorescence Microscopy. Aquatic Microbial Ecology 2006; 43: 223‒231.

27. Herlemann DPR, Labrenz M, Jürgens K, Bertilsson S, Waniek JJ, Andersson AF. Transitions in bacterial communities along the 2000 km salinity gradient of the Baltic Sea. ISME J 2011; 5: 1571‒1579.

28. Tanabe AS. Claident v0.2.2019.05.10. 2019.

29. R Core Team. R: A Language and Environment for Statistical Computing. 2018. Vienna, Austria.

30. Callahan BJ, McMurdie PJ, Rosen MJ, Han AW, Johnson AJA, Holmes SP. DADA2: High-resolution sample inference from Illumina amplicon data. Nat Methods 2016; 13: 581‒583.

31. Ludwig W, Strunk O, Westram R, Richter L, Meier H, Yadhukumar, et al. ARB: a software environment for sequence data. Nucleic Acids Res 2004; 32: 1363‒ 1371.

32. Newton RJ, Jones SE, Eiler A, McMahon KD, Bertilsson S. A Guide to the Natural History of Freshwater Lake Bacteria. Microbiology and Molecular Biology Reviews 2011; 75: 14‒49.

33. Tanabe AS, Toju H. Two New Computational Methods for Universal DNA Barcoding: A Benchmark Using Barcode Sequences of Bacteria, Archaea, Animals, Fungi, and Land Plants. PLoS One 2013; 8: e76910.

34. John SG, Mendez CB, Deng L, Poulos B, Kauffman AKM, Kern S, et al. A simple and efficient method for concentration of ocean viruses by chemical flocculation. Environ Microbiol Rep 2011; 3: 195‒202.

35. Thurber R v, Haynes M, Breitbart M, Wegley L, Rohwer F. Laboratory procedures to generate viral metagenomes. Nat Protoc 2009; 4: 470‒483.

36. Nishimura Y, Watai H, Honda T, Mihara T, Omae K, Roux S, et al. Environmental Viral Genomes Shed New Light on Virus-Host Interactions in the Ocean. mSphere 2017; 2: e00359–16.

37. Hyatt D, Chen G, LoCascio PF, Land ML, Larimer FW, Hauser LJ. Prodigal: prokaryotic gene recognition and translation initiation site identification. BMC Bioinformatics 2010; 11.

38. Edwards RA, McNair K, Faust K, Raes J, Dutilh BE. Computational approaches to predict bacteriophage‒ host relationships. FEMS Microbiol Rev 2016; 40: 258‒272.

39. Kavagutti VS, Andrei A-Ş, Mehrshad M, Salcher MM, Ghai R. Phage-centric ecological interactions in aquatic ecosystems revealed through ultra-deep metagenomics. Microbiome 2019; 7: 135.

40. Chafee M, Fernàndez-Guerra A, Buttigieg PL, Gerdts G, Eren AM, Teeling H, et al. Recurrent patterns of microdiversity in a temperate coastal marine environment. ISME Journal 2018; 12: 237‒ 252.

41. Xia LC, Steele JA, Cram JA, Cardon ZG, Simmons SL, Vallino JJ, et al. Extended local similarity analysis (eLSA) of microbial community and other time series data with replicates. BMC Syst Biol 2011; 5.

42. Xia LC, Ai D, Cram J, Fuhrman JA, Sun F. Efficient statistical significance approximation for local similarity analysis of high-throughput time series data. Bioinformatics 2013; 29: 230‒237.

43. Wickham H, Averick M, Bryan J, Chang W, McGowan L, François R, et al. Welcome to the Tidyverse. J Open Source Softw 2019; 4: 1686.

44. Wickham H. ggplot2: Elegant Graphics for Data Analysis. Springer-Verlag New York 2016.

45. Yamada K, Yamamoto H, Hichiri S, Okamoto T, Hayakawa K. First observation of incomplete vertical circulation in Lake Biwa. Limnology (Tokyo) 2021; 22: 179‒185.

46. Pradeep Ram AS, Nishimura Y, Tomaru Y, Nagasaki K, Nagata T. Seasonal variation in viral-induced mortality of bacterioplankton in the water column of a large mesotrophic lake (Lake Biwa, Japan). Aquatic Microbial Ecology 2010; 58: 249‒259.

47. Arnds J, Knittel K, Buck U, Winkel M, Amann R. Development of a 16S rRNA-targeted probe set for Verrucomicrobia and its application for fluorescence in situ hybridization in a humic lake. Syst Appl Microbiol 2010; 33: 139‒148.

48. He S, Stevens SLR, Chan L-K, Bertilsson S, Glavina del Rio T, Tringe SG, et al. Ecophysiology of Freshwater Verrucomicrobia Inferred from Metagenome-Assembled Genomes. mSphere 2017; 2.

49. Goldfarb KC, Karaoz U, Hanson CA, Santee CA, Bradford MA, Treseder KK, et al. Differential growth responses of soil bacterial taxa to carbon substrates of varying chemical recalcitrance. Front Microbiol 2011; 2.

50. Takasu H, Ushio M, LeClair J, Nakano S. High contribution of Synechococcus to phytoplankton biomass in the aphotic hypolimnion in a deep freshwater lake (Lake Biwa, Japan). Aquatic Microbial Ecology 2015; 75: 69‒79.

51. Agawin N, Duarte C, Agustí S. Growth and abundance of Synechococcus sp. in a Mediterranean Bay:seasonality and relationship with temperature. Mar Ecol Prog Ser 1998; 170: 45‒53.

52. Kolmonen E, Sivonen K, Rapala J, Haukka K. Diversity of cyanobacteria and heterotrophic bacteria in cyanobacterial blooms in Lake Joutikas, Finland. Aquatic Microbial Ecology 2004; 36: 201‒211.

53. Thottathil SD, Hayakawa K, Hodoki Y, Yoshimizu C, Kobayashi Y, Nakano S. Biogeochemical control on fluorescent dissolved organic matter dynamics in a large freshwater lake (Lake Biwa, Japan). Limnol Oceanogr 2013; 58: 2262‒2278.

54. Tinta T, Christiansen LS, Konrad A, Liberles DA, Turk V, Munch-Petersen B, et al. Deoxyribonucleoside kinases in two aquatic bacteria with high specificity for thymidine and deoxyadenosine. FEMS Microbiol Lett 2012; 331: 120‒127.

55. Tada Y, Grossart H-P. Community shifts of actively growing lake bacteria after N-acetyl-glucosamine addition: improving the BrdU-FACS method. ISME J 2014; 8: 441‒454.

56. Nakano S, Koitabashi T, Ueda T. Seasonal changes in abundance of heterotrophic nanoflagellates and their consumption of bacteria in Lake Biwa with special reference to trophic interactions with Daphnia galeata. Fundamental and Applied Limnology 1998; 142: 21‒34.

57. Luo E, Leu AO, Eppley JM, Karl DM, DeLong EF. Diversity and origins of bacterial and archaeal viruses on sinking particles reaching the abyssal ocean. ISME J 2022.

58. Yamada Y, Tomaru Y, Fukuda H, Nagata T. Aggregate formation during the viral lysis of a marine diatom. Front Mar Sci 2018; 5: 1‒7.

59. Paul JH. Prophages in marine bacteria: dangerous molecular time bombs or the key to survival in the seas? ISME J 2008; 2: 579‒589.

60. Jiang SC, Paul JH. Significance of Lysogeny in the Marine Environment: Studies with Isolates and a Model of Lysogenic Phage Production. Microb Ecol 1998; 35: 235‒243.

61. Knowles B, Silveira CB, Bailey BA, Barott K, Cantu VA, Cobián-Güemes AG, et al. Lytic to temperate switching of viral communities. Nature 2016; 531: 466‒470.

62. Ikeda S, Ichise S, Furuta S, Urabe J. Long-term Changes in Seasonality of Phytoplankton Community in North Basin of Lake Biwa: a Comparison with the Plankton Ecology Group (PEG) Model. Journal of Japan Society on Water Environment 2018; 41: 115‒ 122.

63. Kishimoto N, Yamamoto C, Suzuki K, Ichise S. Does a Decrease in Chlorophyll a Concentration in Lake Biwa Mean a Decrease in Primary Productivity by Phytoplankton? J Water Environ Technol 2015; 13: 1‒14.

64. Wiggins BA, Alexander M. Minimum bacterial density for bacteriophage replication: implications for significance of bacteriophages in natural ecosystems. Appl Environ Microbiol 1985; 49: 19‒23.

