## Supplementary material for "Identification of key virus–prokaryote infection pairs that contribute to viral shunt in a freshwater lake": Fig. S1, Fig. S2, Fig. S3, Fig. S4, Fig. S5, Table S1, Table S2, Table S3, Table S4, Table S5, Table S6

This PDF file includes:

Materials and methods

Fig. S1
Fig. S2
Fig. S3
Fig. S4
Fig. S5

Table S1
Table S2
Table S3
Table S4
Table S5
Table S6

#### Materials and methods

##### Genome assembly and gene prediction and annotation

Low-quality raw reads were removed using Trimmomatic v0.39 with default settings [1]. Thereafter, raw reads were assembled using SPAdes v. 3.13.1 with the "—careful" option [2, 3]. All 27 samples were assembled using SPAdes using the same options. Only contigs longer than 10 kbp (29 730 contigs) were used for further analysis. VirSorter removes non-viral contigs using the "--virome" option [4]. Contigs classified as VirSorter category 1-3 ("most confident," "likely," and "possible" prediction, [4]) were considered viral contigs and used for further analysis (Table S3, 24 967 contigs). These viral contigs were clustered using all-vs.-all blastn at >95% nucleotide identity across >95% of their length to remove redundancy [5]. The longest contig in each cluster was selected as the representative contig. The complete genome (i.e., circular genome) was determined using ccfind with default settings [3]. Open reading frames were predicted using Prodigal v. 2.6.3 with the "-p meta" option [6]. The gene function of the predicted open reading frames was annotated according to a previous study (see "Workflow for gene functional annotation" in the supplementary information [7]).

##### Host prediction

Lake Biwa virus (LBV) hosts were assigned using multiple methods [8]. Actinobacteria and Cyanobacteria were predicted to be hosts of an LBV when host-specific genes were found in a phage genome/contig (i.e., *whiB* for Actinobacteria [9] and a photosystem gene for Cyanobacteria [10]). CRISPR spacers and tRNAs in prokaryotic genomes obtained from the NCBI database (search condition: "(BCT) AND (WGS) AND ((Lake freshwater) OR (freshwater))") and a previous study [7] were detected using metaCRT [11] and ARAGORN1.2.36 [12], respectively. LBV sequences were compared to the CRISPR spacers using BLASTn with a threshold of 100% identity and to the tRNAs using BLASTn with thresholds > 97% identity, alignment length ≥ 30 bp, and < $10^{-5}$ e-value. LBV genomes were also directly compared with prokaryotic genomes using BLASTn with thresholds > 97% identity, alignment length ≥ 30 bp, and < $10^{-5}$ e-value. The prokaryotic genomes used for this prediction (using CRISPR, tRNA, and direct comparison) were obtained from NCBI RefSeq (downloaded in June 2020) and Lake Biwa metagenome-assembled genomes [7]. Other methods for host prediction were “A close relative infected a known host” and “>80% of the annotated bacterial genes were taxonomically related to a single taxon,” as described earlier [7].

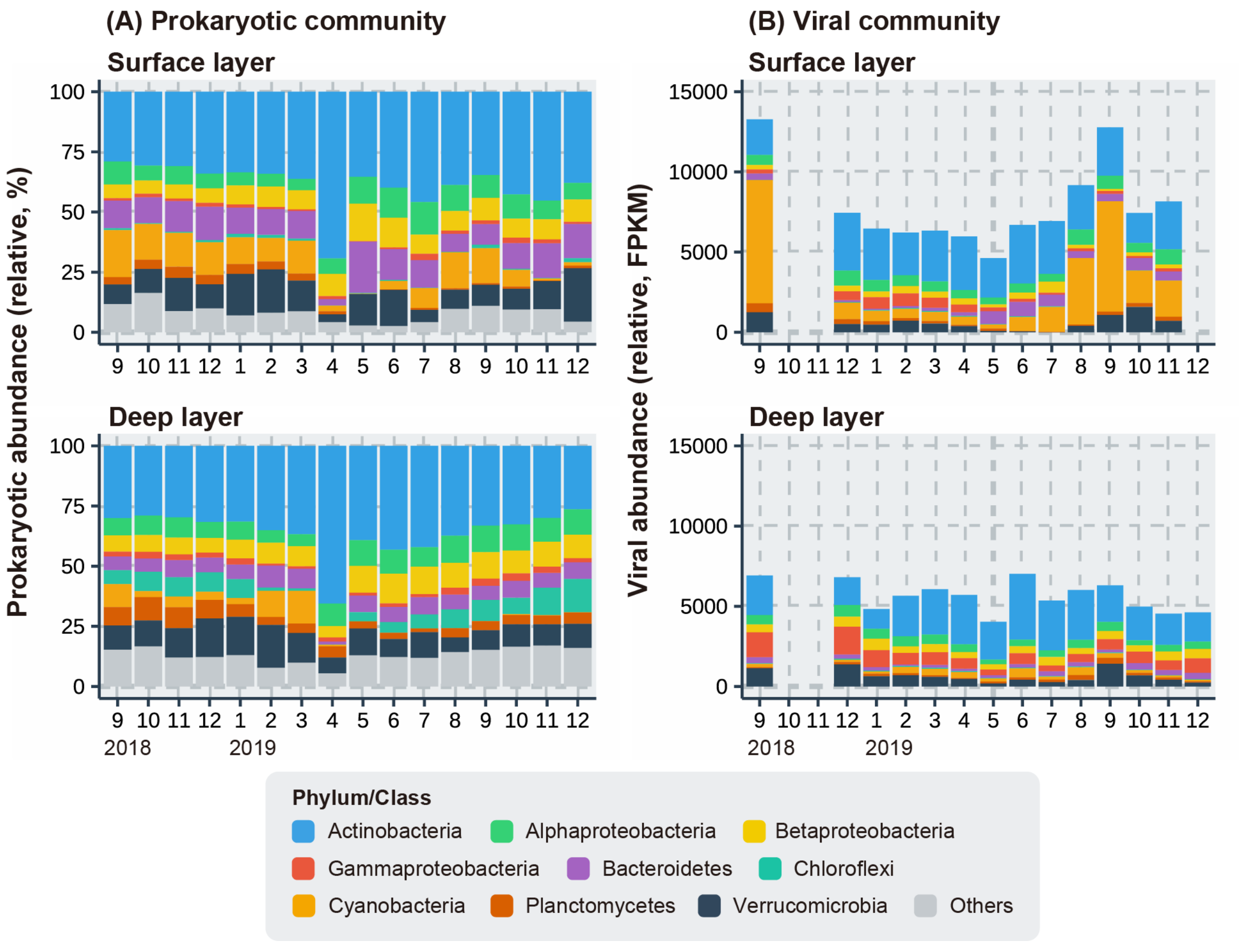


**Fig. S1.** Seasonal variation in the (A) prokaryotic community, shown as a percentage, and (B) viral community, shown as FPKM, in the surface and deep layers during the study period. Viruses whose hosts were not predicted or the hosts shown in the legend that were not predicted were excluded from the figures. Prokaryotes were classified by their phylum/class and viruses based on their predicted host phylum/class.

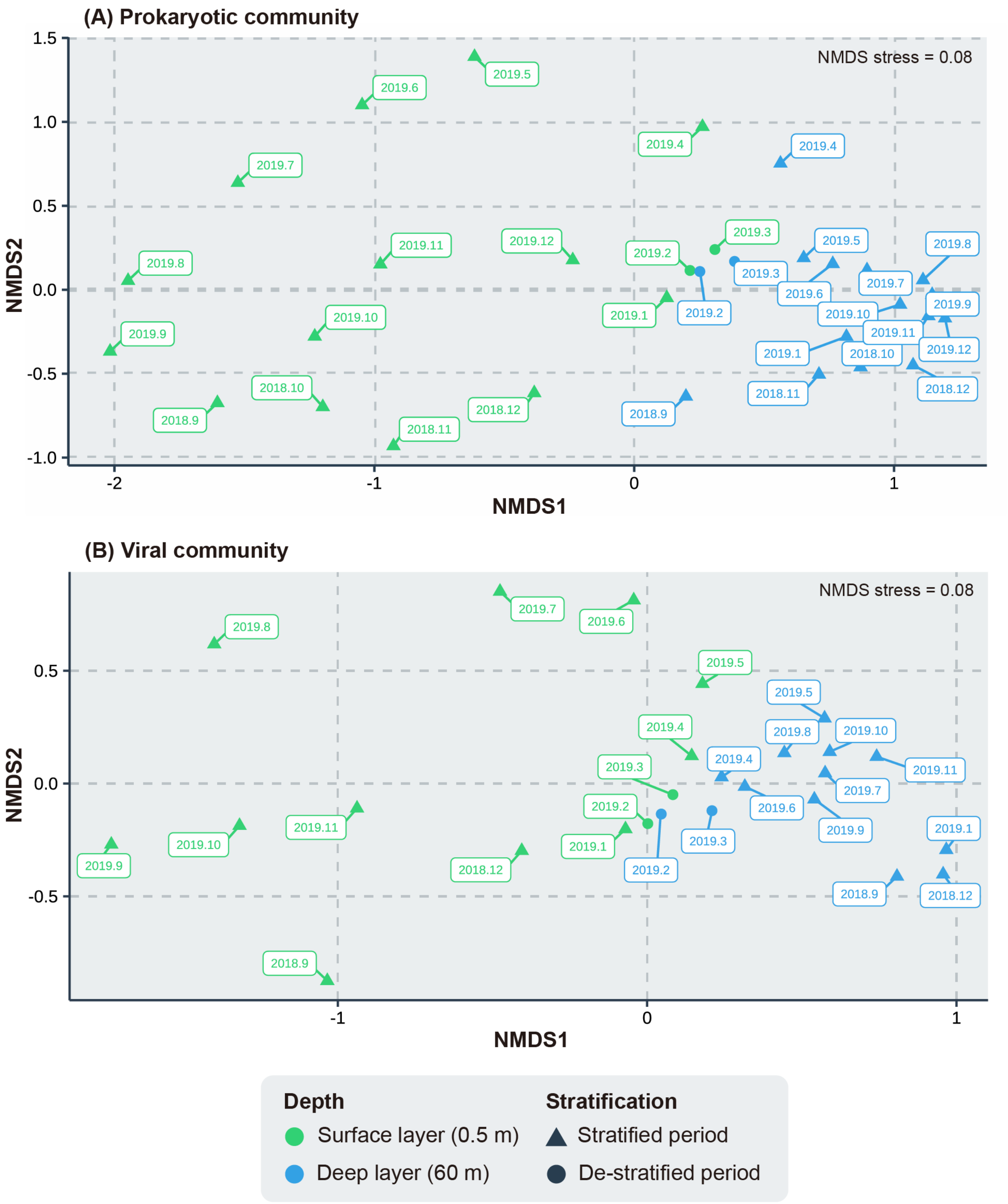


**Fig. S2.** Non-metric Multi-dimensional Scaling (NMDS) plots of (A) prokaryotic and (B) viral communities in the surface and deep layers during the study period

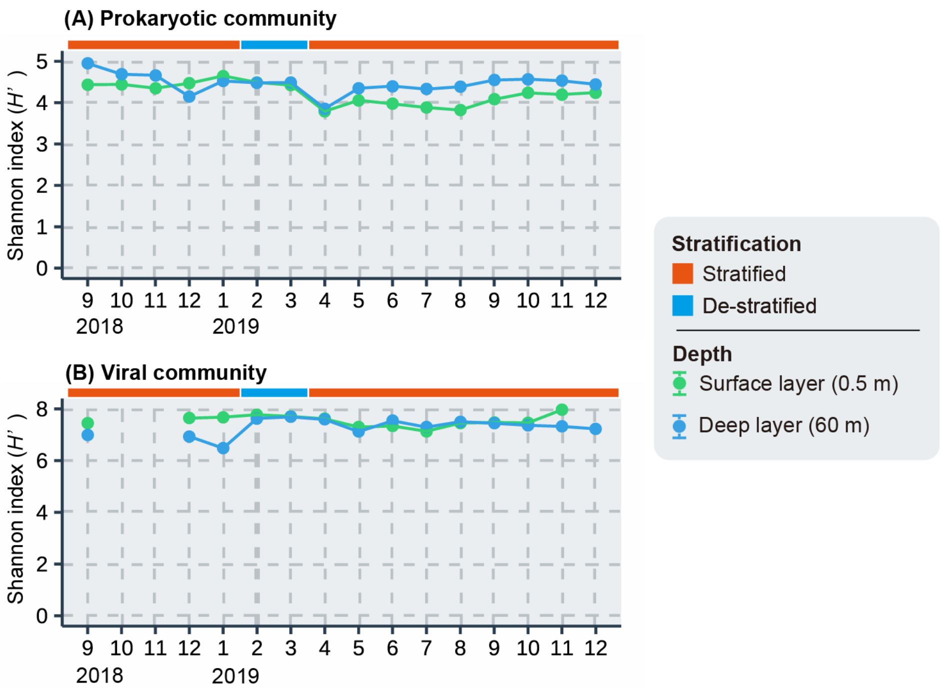


**Fig. S3.** Seasonal variation in the alpha diversity of (A) prokaryotic and (B) viral communities in the surface and deep layers

#### Actinobacteria and its phages in the surface layer (20 ASVs)

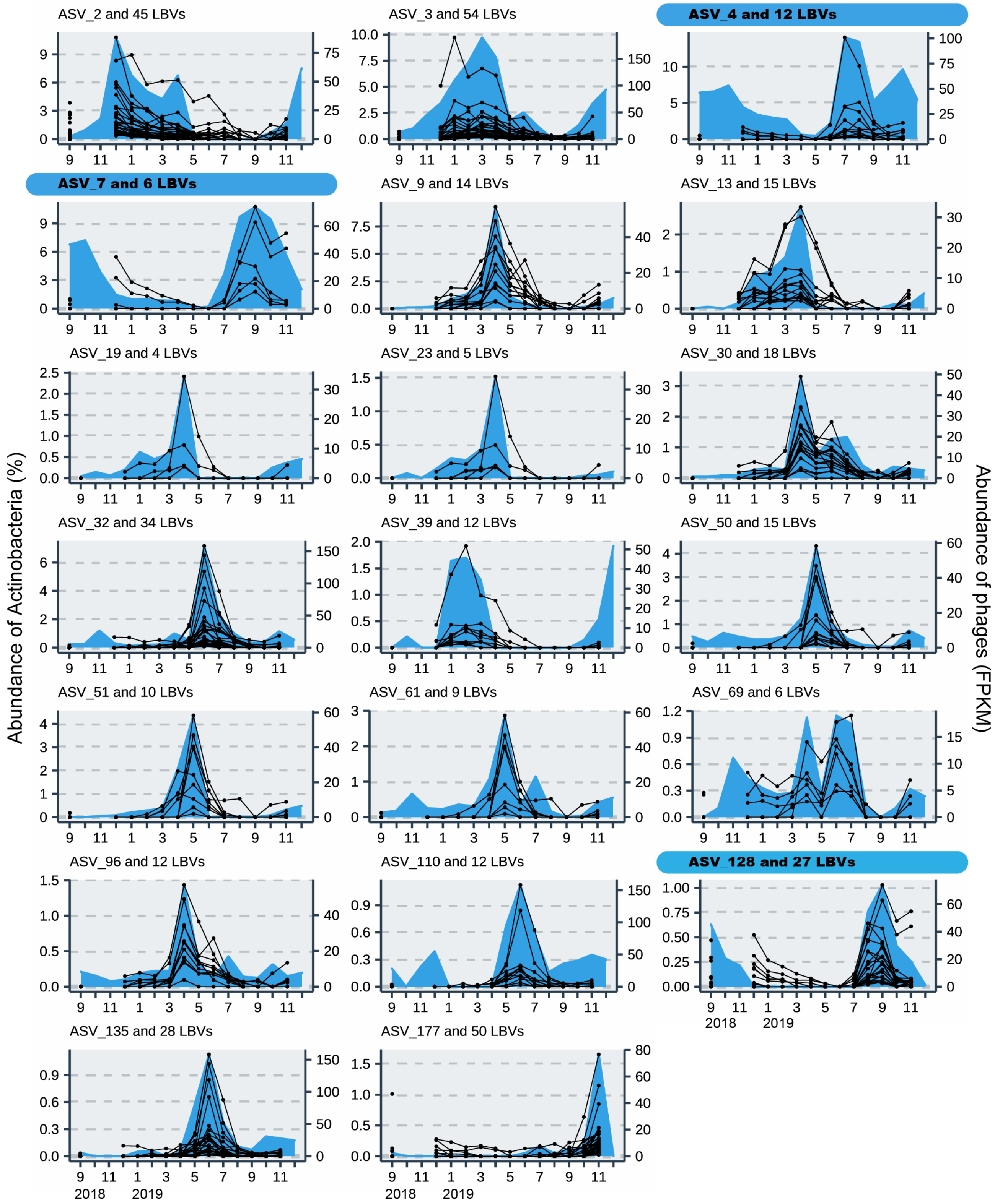


**Fig. S4.** Seasonal abundance of prokaryotic species (ASV) and viruses (LBV) co-occurring with the species in the surface layer. The subtitle of each panel indicates the ID of the ASV and the number of LBVs that co-occur with that ASV. Shaded subtitles indicate ASVs estimated to contribute to prokaryotic production in summer (July to

September 2019) and co-occurring LBVs. ASVs indicate amplicon sequence variants, and LBVs indicate Lake Biwa viruses.

#### Alphaproteobacteria and its phages in the surface layer (6 ASVs)

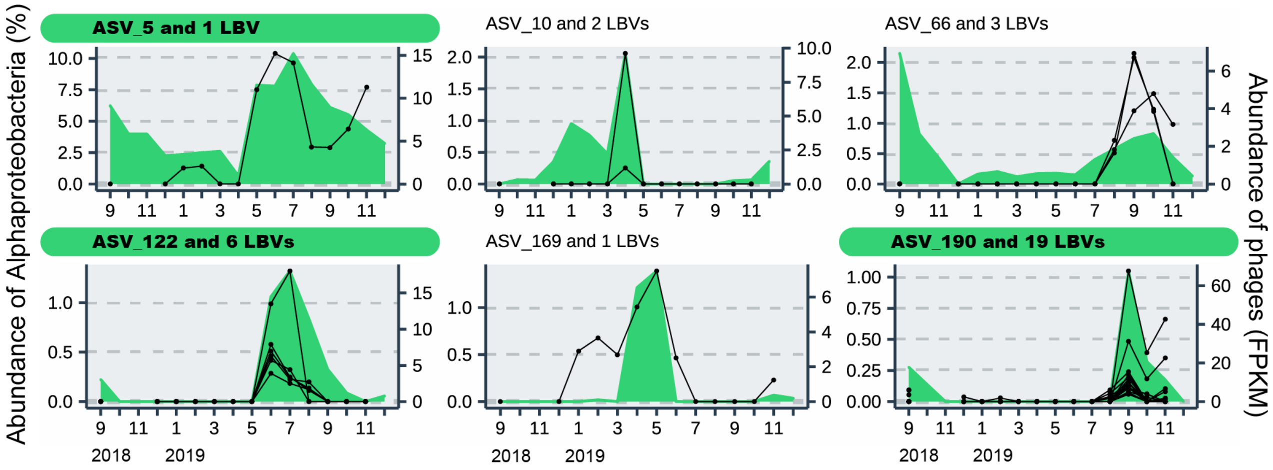


#### Betaproteobacteria and its phages in the surface layer (7 ASVs)

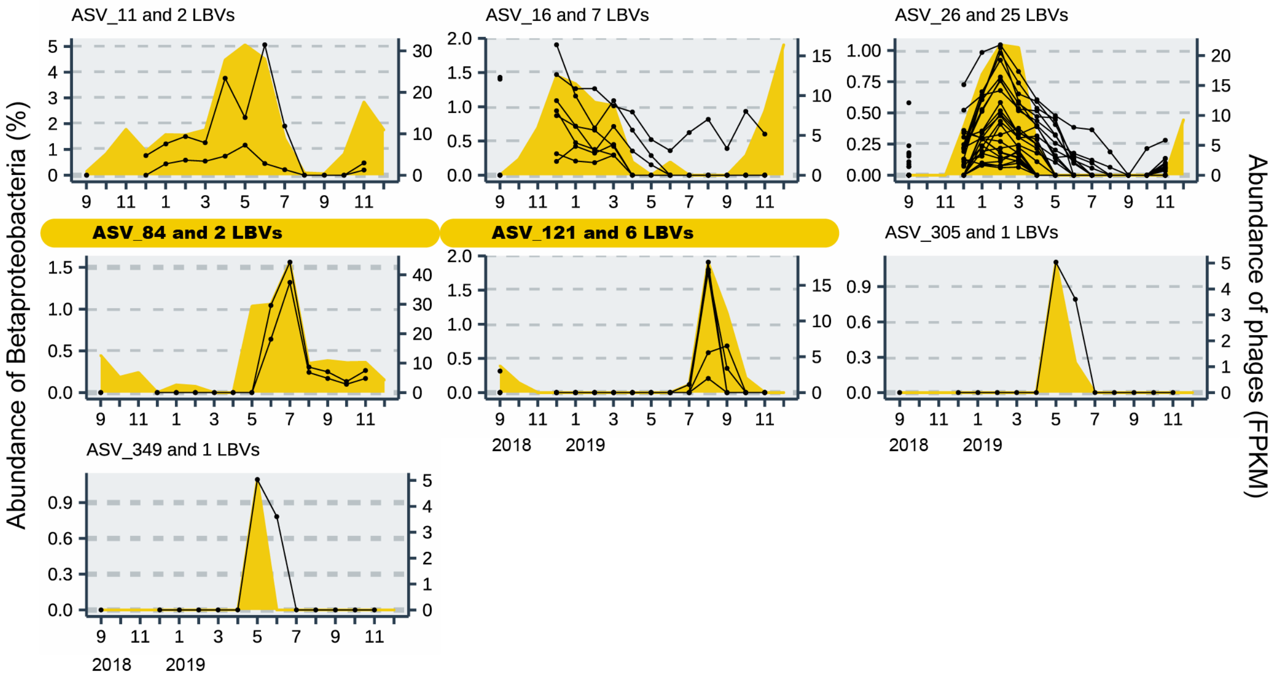


**Fig. S4 (continued)**

### Bacteroidetes and its phages in the surface layer (20 ASVs)

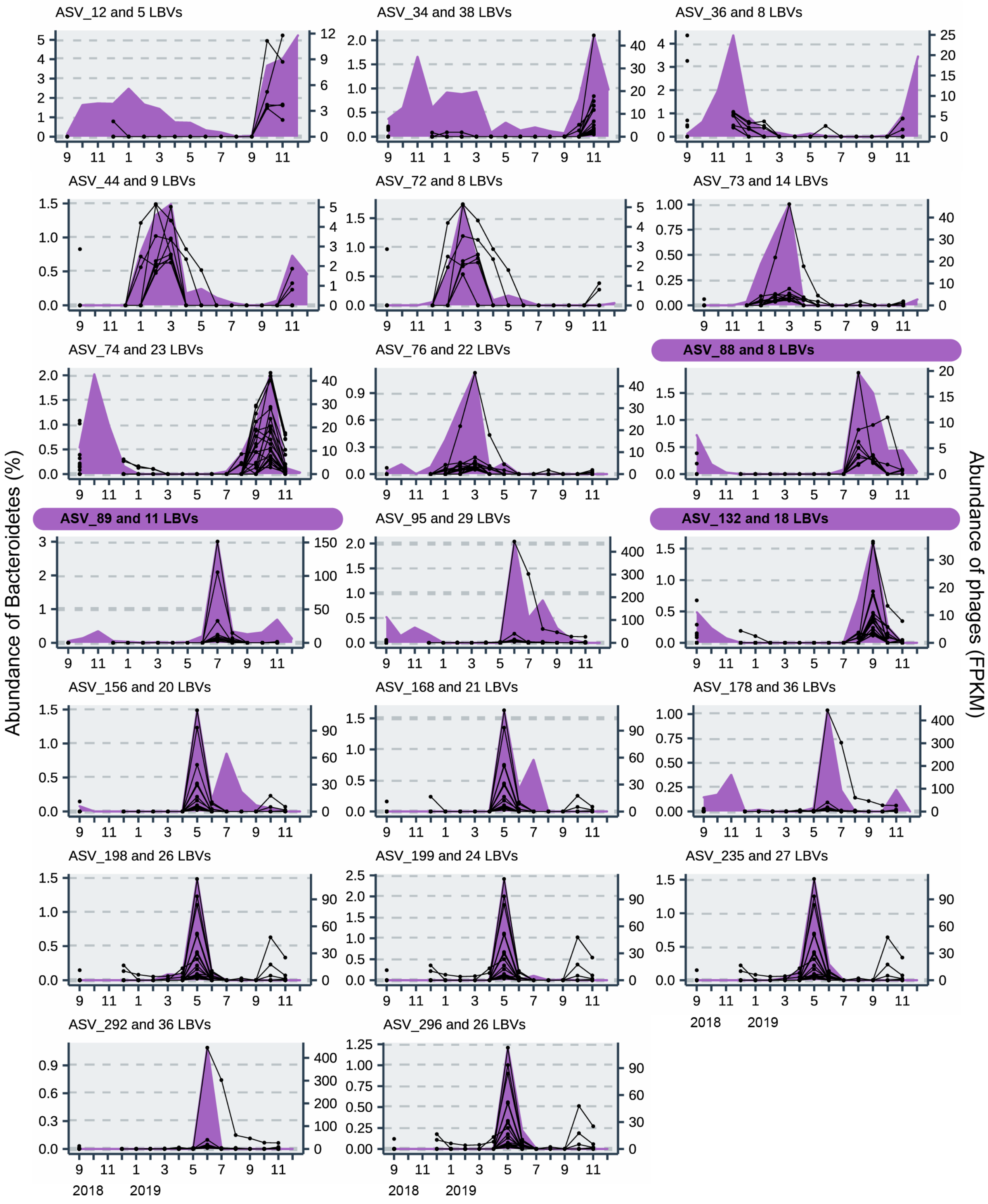


**Fig. S4 (continued)**

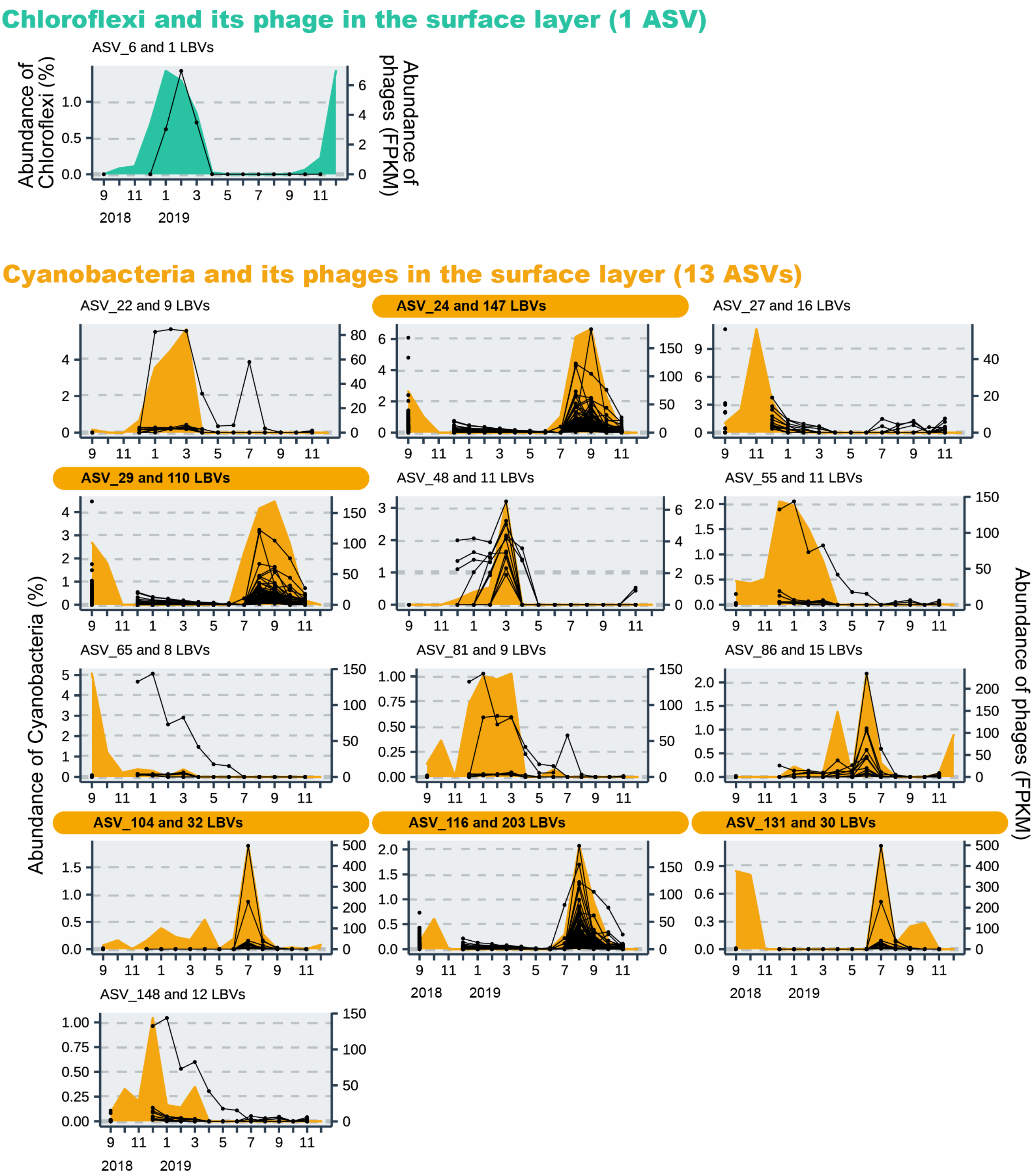


**Fig. S4 (continued)**

#### Verrucomicrobia and its phages in the surface layer (15 ASVs)

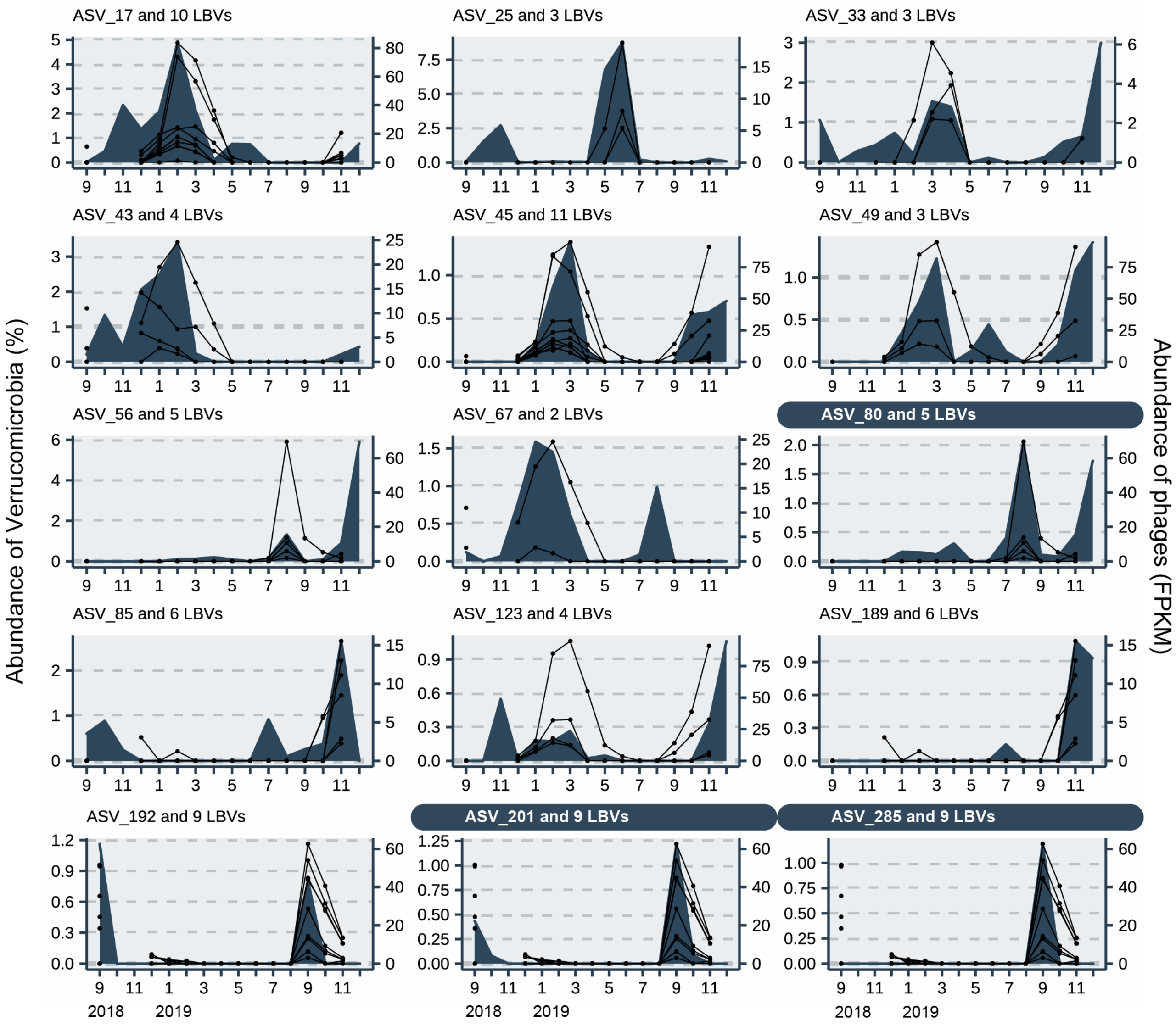


**Fig. S4 (continued)**

#### Actinobacteria and its phages in the deep layer (8 ASVs)

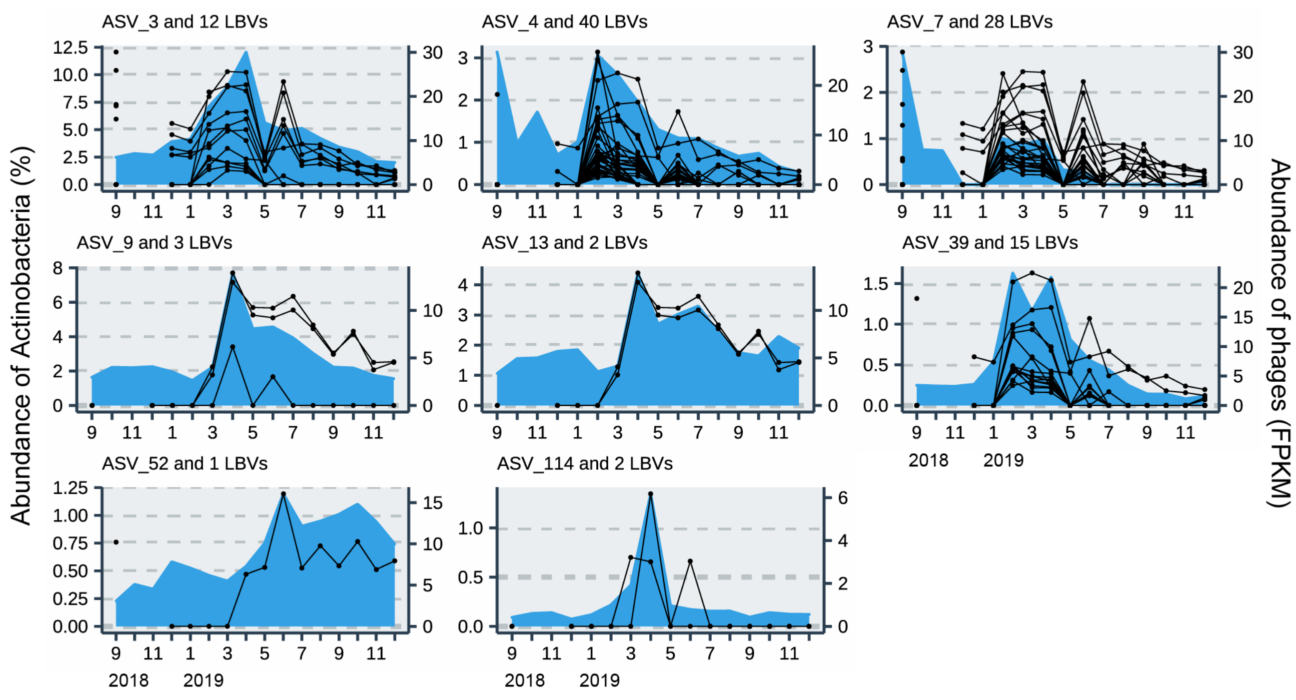


#### Betaproteobacteria and its phages in the deep layer (2 ASVs)

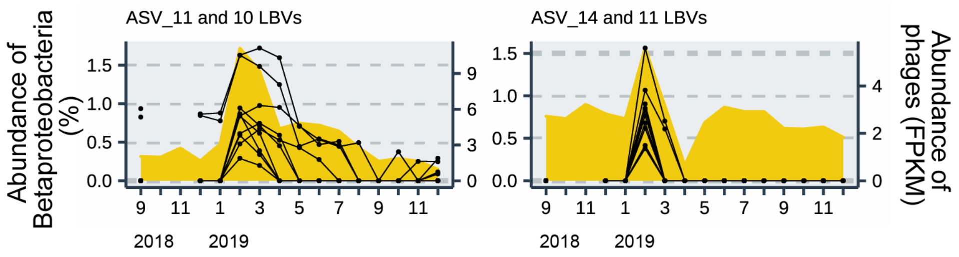


#### Bacteroidetes and its phage in the deep layer (1 ASV)

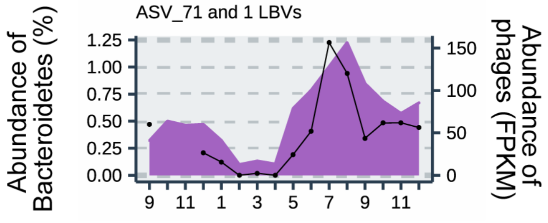


**Fig. S5.** Seasonal abundance of prokaryotic species (ASV) and viruses (LBV) co-occurring with the species in the deep layers. The subtitle of each panel indicates the ID of the ASV and the number of LBVs that co-occurred with the ASV. ASVs indicate amplicon sequence variants, and LBVs indicate Lake Biwa viruses.

#### Cyanobacteria and its phages in the deep layer (6 ASVs)

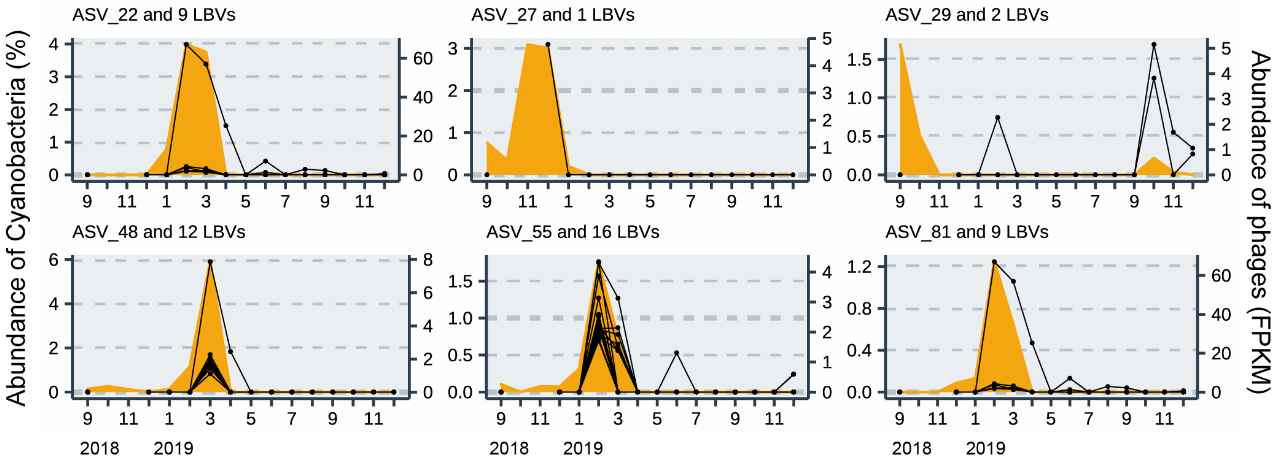


#### Planctomycetes and its phage in the deep layer (1 ASV)

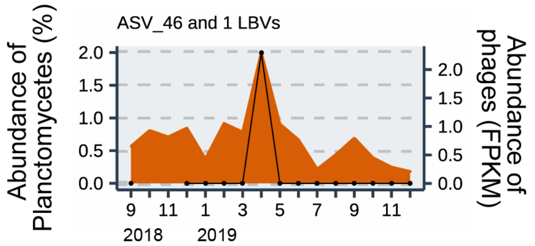


#### Verrucomicrobia and its phages in the deep layer (8 ASVs)

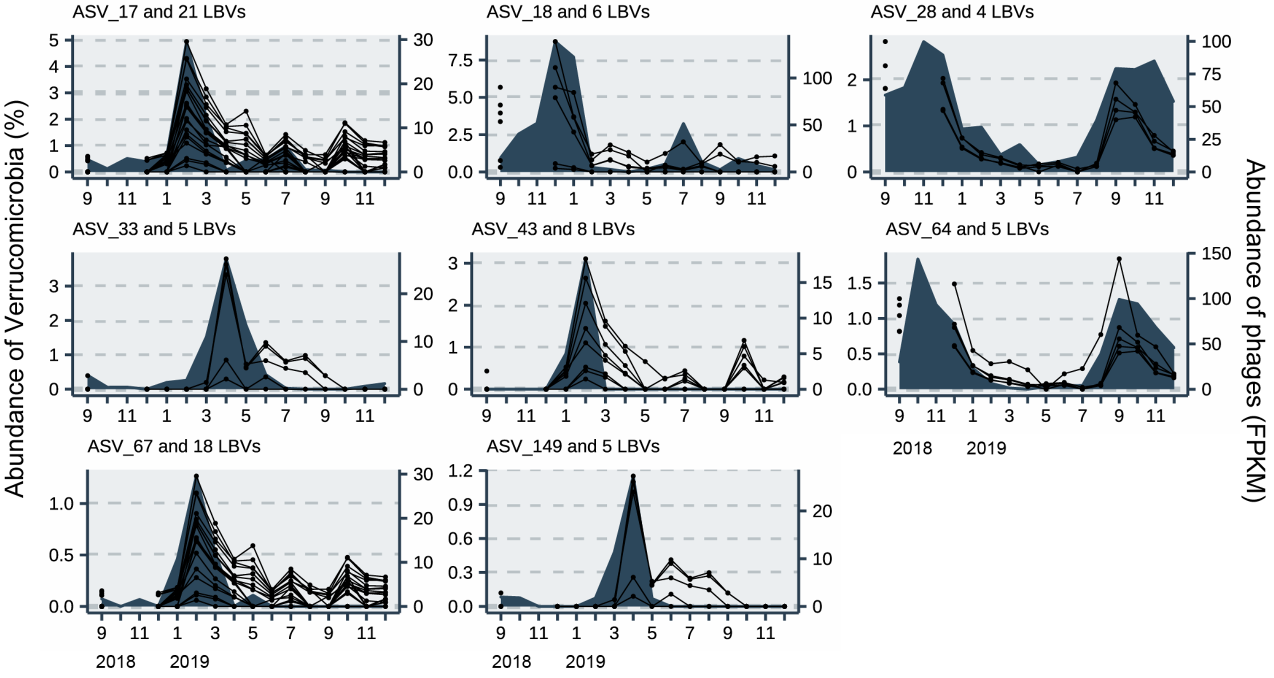


**Fig. S5 (continued)**

**Table S1.** Basic information on sample collection

| Parameters | 2018<br>9 | 10 | 11 | 12 | 2019<br>1 | 2 | 3 | 4 | 5 | 6 | 7 | 8 | 9 | 10 | 11 | 12 |
|---|---|---|---|---|---|---|---|---|---|---|---|---|---|---|---|---|
| Environmental parameters [a] | ■ | ■ | ■ | ■ | ■ | ■ | ■ | ■ | ■ | ■ | ■ | ■ | ■ | ■ | ■ | ■ |
| Prokaryotic community | ■ | ■ | ■ | ■ | ■ | ■ | ■ | ■ | ■ | ■ | ■ | ■ | ■ | ■ | ■ | ■ |
| Viral community | ■ | | | ■ | ■ | ■ | ■ | ■ | ■ | ■ | ■ | ■ | ■ | ■ | ■ | ■ b |
| Prokaryotic production | | | | | | | | | ■ | ■ | ■ | ■ | ■ | ■ | ■ | |

[a] The environmental parameters included water temperature and dissolved oxygen concentration.

[b] Sample collected only from the surface layer

**Table S2.** Samples and sequence reads of 16S rRNA used in this study

(A) Surface layer (0. 5 m)

| Date (YYYY/MM/DD) | Raw reads | Denoised forward reads | Denoised reverse reads | Merged reads | Non-chimera reads |
|---|---|---|---|---|---|
| 2018/9/18 | 66,263 | 61,314 | 64,138 | 49,252 | 44,193 |
| 2018/10/15 | 54,093 | 49,913 | 51,933 | 39,299 | 34,984 |
| 2018/11/19 | 65,596 | 60,555 | 63,563 | 49,327 | 44,140 |
| 2018/12/17 | 51,274 | 47,986 | 49,516 | 40,490 | 39,294 |
| 2019/1/18 | 64,285 | 59,638 | 62,524 | 51,154 | 48,132 |
| 2019/2/18 | 82,194 | 75,455 | 80,009 | 64,457 | 59,228 |
| 2019/3/18 | 71,617 | 67,273 | 70,059 | 58,926 | 54,398 |
| 2019/4/22 | 62,796 | 60,706 | 61,879 | 54,397 | 49,795 |
| 2019/5/21 | 53,708 | 50,988 | 52,617 | 43,455 | 40,458 |
| 2019/6/17 | 75,177 | 71,639 | 73,958 | 63,110 | 57,431 |
| 2019/7/16 | 67,905 | 64,768 | 66,853 | 57,150 | 51,884 |
| 2019/8/19 | 77,356 | 72,883 | 75,896 | 63,498 | 57,396 |
| 2019/9/17 | 71,530 | 67,413 | 69,625 | 58,413 | 54,047 |
| 2019/10/21 | 83,464 | 78,503 | 81,315 | 66,550 | 61,117 |
| 2019/11/18 | 70,731 | 67,124 | 69,176 | 57,218 | 52,691 |
| 2019/12/16 | 62,948 | 58,412 | 61,551 | 50,428 | 46,252 |
| Total number | 1,080,937 | 1,014,570 | 1,054,612 | 867,124 | 795,440 |

(B) Deep layer (60 m)

| Date (YYYY/MM/DD) | Raw reads | Denoised forward reads | Denoised reverse reads | Merged reads | Non-chimera reads |
|---|---|---|---|---|---|
| 2018/9/18 | 54,077 | 49,763 | 51,756 | 40,248 | 38,419 |
| 2018/10/15 | 56,743 | 51,368 | 54,060 | 40,208 | 36,769 |
| 2018/11/19 | 73,602 | 67,426 | 70,608 | 53,929 | 48,903 |
| 2018/12/17 | 54,378 | 50,718 | 51,411 | 40,999 | 39,579 |
| 2019/1/18 | 60,488 | 56,201 | 58,292 | 48,291 | 45,913 |
| 2019/2/18 | 71,540 | 66,234 | 69,764 | 57,135 | 53,376 |
| 2019/3/18 | 77,741 | 72,865 | 75,768 | 63,577 | 59,097 |
| 2019/4/22 | 53,161 | 50,953 | 52,085 | 45,090 | 40,339 |
| 2019/5/21 | 59,595 | 56,504 | 58,313 | 48,879 | 44,861 |
| 2019/6/17 | 54,417 | 51,127 | 52,971 | 43,092 | 38,970 |
| 2019/7/16 | 58,575 | 55,112 | 56,870 | 47,139 | 43,456 |
| 2019/8/19 | 49,232 | 46,125 | 47,841 | 38,241 | 35,334 |
| 2019/9/17 | 66,326 | 61,028 | 64,541 | 52,117 | 48,168 |
| 2019/10/21 | 55,075 | 50,513 | 52,968 | 41,841 | 39,446 |
| 2019/11/18 | 67,904 | 62,841 | 65,730 | 54,151 | 50,956 |
| 2019/12/16 | 51,814 | 47,823 | 50,089 | 40,680 | 38,701 |
| Total number | 964,668 | 896,601 | 933,067 | 755,617 | 702,287 |

**Table S3.** Virome samples and sequence reads used in this study

| Date (YYYY/MM/DD) | The surface layer (0.5 m) | | | | |
|---|---|---|---|---|---|
| | Raw reads | Quality controlled reads | Contigs (>10 kbp) | Contigs (>10 kbp & VirSorter) | Complete genomes |
| 2018/9/18 | 7,323,537 | 7,019,303 | 687 | 637 | 20 |
| 2018/10/15 | | | | | |
| 2018/11/19 | | | | | |
| 2018/12/17 | 5,153,761 | 4,921,629 | 791 | 725 | 28 |
| 2019/1/18 | 4,847,177 | 4,612,254 | 621 | 548 | 10 |
| 2019/2/18 | 5,583,923 | 5,406,611 | 836 | 753 | 15 |
| 2019/3/18 | 3,689,875 | 3,559,368 | 725 | 626 | 24 |
| 2019/4/22 | 3,641,313 | 3,483,517 | 832 | 763 | 25 |
| 2019/5/21 | 3,669,706 | 3,369,218 | 766 | 691 | 4 |
| 2019/6/17 | 4,243,551 | 4,044,155 | 672 | 598 | 15 |
| 2019/7/16 | 6,488,878 | 6,173,660 | 653 | 586 | 13 |
| 2019/8/19 | 6,218,320 | 5,948,181 | 1,078 | 901 | 20 |
| 2019/9/17 | 4,318,398 | 4,110,974 | 649 | 597 | 8 |
| 2019/10/21 | 3,162,795 | 2,969,758 | 787 | 704 | 26 |
| 2019/11/18 | 11,943,151 | 11,505,833 | 1,773 | 1,602 | 79 |
| 2019/12/16 | | | | | |
| **Total number** | **70,284,385** | **67,124,461** | **10,870** | **9,731** | **287** |

| Date (YYYY/MM/DD) | The Deep layer (60 m) | | | | |
|---|---|---|---|---|---|
| | Raw reads | Quality Controlled reads | Contigs (>10 kbp) | Contigs (>10 kbp & VirSorter) | Complete genomes |
| 2018/9/18 | 4,865,998 | 4,683,162 | 256 | 221 | 16 |
| 2018/10/15 | | | | | |
| 2018/11/19 | | | | | |
| 2018/12/17 | 4,745,284 | 4,546,583 | 303 | 269 | 7 |
| 2019/1/18 | 5,123,480 | 4,921,742 | 282 | 242 | 12 |
| 2019/2/18 | 3,722,244 | 3,586,393 | 899 | 818 | 25 |
| 2019/3/18 | 4,772,796 | 4,612,203 | 713 | 626 | 14 |
| 2019/4/22 | 3,340,409 | 3,228,368 | 842 | 772 | 28 |
| 2019/5/21 | 2,539,636 | 2,340,002 | 866 | 606 | 17 |
| 2019/6/17 | 6,384,547 | 6,062,560 | 580 | 515 | 11 |
| 2019/7/16 | 4,674,055 | 4,466,927 | 743 | 673 | 17 |
| 2019/8/19 | 4,133,169 | 3,954,145 | 702 | 432 | 11 |
| 2019/9/17 | 4,208,646 | 3,996,723 | 750 | 657 | 18 |
| 2019/10/21 | 4,929,853 | 4,686,369 | 902 | 609 | 20 |
| 2019/11/18 | 4,504,833 | 4,335,050 | 782 | 684 | 43 |
| 2019/12/16 | 12,520,512 | 11,952,128 | 1,221 | 874 | 38 |
| **Total number** | **70,465,462** | **67,372,355** | **9,841** | **7,998** | **277** |

| All samples (all months and depths) | | |
|---|---|---|
| Contigs (>10 kbp) | Contigs (>10 kbp & VirSorter) | Complete genomes |
| 9,019 | 7,238 | 254 |

**Table S4.** The numbers of amplicon sequence variants and Lake Biwa viruses in the hosts predicted in this study

| Phylum/class | Prokaryotes | | Viruses whose hosts were predicted | |
|---|---|---|---|---|
| | # | Proportion (%) | # | Proportion (%) |
| Actinobacteria | 118 | 7.3 | 670 | 24.1 |
| Alphaproteobacteria | 230 | 14.3 | 274 | 9.9 |
| Betaproteobacteria | 180 | 11.2 | 48 | 1.7 |
| Gammaproteobacteria | 135 | 8.4 | 243 | 8.7 |
| Bacteroidetes | 261 | 16.2 | 322 | 11.6 |
| Chloroflexi | 12 | 0.7 | 13 | 0.5 |
| Cyanobacteria | 89 | 5.5 | 784 | 28.7 |
| Planctomycetes | 86 | 5.3 | 83 | 3.0 |
| Verrucomicrobia | 154 | 9.6 | 99 | 3.6 |
| Others | 343 | 21.3 | 244 | 8.8 |
| Total | 1,608 | 100 | 2,780 | 100 |

**Table S5.** Statistical significance of the time separation dissimilarity difference between the surface and deep layers. The Mann–Whitney *U* test and Student's *t*-test were used to assess prokaryotic and viral communities, respectively.

| | *p*-value | |
|---|---|---|
| **Sample separation (months)** | **Prokaryotes** | **Viruses** |
| 1 | (0.0037) ** | (0.16) NS |
| 2 | $(9.7 \times 10^{-6})$ *** | (0.0079) ** |
| 3 | $(1.3 \times 10^{-6})$ *** | $(2.0 \times 10^{-4})$ *** |
| 4 | $(1.4 \times 10^{-5})$ *** | $(9.4 \times 10^{-6})$ *** |
| 5 | $(2.8 \times 10^{-6})$ *** | $(3.5 \times 10^{-5})$ *** |
| 6 | $(1.1 \times 10^{-5})$ *** | $(1.0 \times 10^{-6})$ *** |
| 7 | $(4.1 \times 10^{-5})$ *** | $(1.8 \times 10^{-5})$ *** |
| 8 | (0.0011) ** | (0.0010) ** |
| 9 | (0.0070) ** | (0.011) * |
| 10 | (0.026) * | (0.16) NS |
| 11 | (0.032) * | NA |
| 12 | (0.34) NS | NA |
| 13 | (0.20) NS | NA |
| 14 | NA | NA |
| 15 | NA | NA |

The original data are shown in Fig. 2C. *$p$ <0.05; **$p$ <0.01; ***$p$ <0.001; NSNo significant difference. NA: Statistical tests (i.e., Mann–Whitney *U* test or Student's *t*-test) were not applied as the sample sizes were less than 3.

**Table S6.** (a) Number of amplicon sequence variants (ASVs) dominant in the middle of the stratified period ($ASV_{dominant}$) and their relative abundance (%) in 2019. (b) The number of lake Biwa viruses (LBV) co-occurring with the $ASV_{dominant}$. * Gemmatimonadetes was excluded from the analysis because its phages could not be predicted.

(a) Prokaryotes ($ASV_{dominant}$)

| Phylum/class | # | Relative abundance (%) July | August | September |
|---|---|---|---|---|
| **Actinobacteria** | 3 | 17.8 | 24.0 | 17.1 |
| **Alphaproteobacteria** | 3 | 11.7 | 8.8 | 7.5 |
| **Betaproteobacteria** | 2 | 1.7 | 2.3 | 1.6 |
| **Bacteroidetes** | 3 | 3.1 | 2.9 | 3.4 |
| **Cyanobacteria** | 5 | 6.6 | 12.7 | 12.3 |
| **Verrucomicrobia** | 3 | 0.4 | 2.1 | 2.5 |
| **Chlorobi** | 1 | 0.13 | 2.6 | 1.6 |
| **Gemmatimonadetes** | 1 | 0.0 | 1.1 | 0.38 |
| **Total** | 21 | 41.3 | 56.3 | 46.2 |

(b) Viruses (LBV)

| Host phylum/class | # | FPKM-based relative abundance (%) July | August | September |
|---|---|---|---|---|
| **Actinobacteria** | 39 | 0.54 | 1.4 | 1.2 |
| **Alphaproteobacteria** | 26 | 0.10 | 0.070 | 0.52 |
| **Betaproteobacteria** | 8 | 0.17 | 0.21 | 0.046 |
| **Bacteroidetes** | 33 | 0.72 | 0.20 | 0.44 |
| **Cyanobacteria** | 276 | 2.8 | 8.7 | 4.8 |
| **Verrucomicrobia** | 14 | 0.0043 | 0.23 | 0.60 |
| **Chlorobi** | 0 | | | |
| **Gemmatimonadetes** | * | | | |
| **Total** | 396 | 4.3 | 10.8 | 7.6 |

**Table S7.** Summary of amplicon sequence variants (ASVs) detected in this study

**Table S8.** Summary of Lake Biwa viruses (LBVs) detected in this study

**Table S9.** Dominance patterns of Lake Biwa viruses (LBVs) ranked within 100 based on fragments per kilobase of contig per million mapped reads (FPKM) abundance in the surface and deep layers. "1" in the month column shows that the LBVs ranked within the 100 of that month. Numbers shown in the "Period (Mo)" column indicate the total dominant duration (months) through this study.

**References**


1. Bolger AM, Lohse M, Usadel B. Trimmomatic: A flexible trimmer for Illumina sequence data. *Bioinformatics* 2014; **30**: 2114–2120.
2. Bankevich A, Nurk S, Antipov D, Gurevich AA, Dvorkin M, Kulikov AS, et al. SPAdes: a new genome assembly algorithm and its applications to single-cell sequencing. *J Comput Biol* 2012; **19**: 455–477.
3. Nishimura Y, Watai H, Honda T, Mihara T, Omae K, Roux S, et al. Environmental Viral Genomes Shed New Light on Virus-Host Interactions in the Ocean. *mSphere* 2017; **2**: e00359-16.
4. Roux S, Enault F, Hurwitz BL, Sullivan MB. VirSorter: mining viral signal from microbial genomic data. *PeerJ* 2015; **3**: e985.
5. Kavagutti VS, Andrei A-Ș, Mehrshad M, Salcher MM, Ghai R. Phage-centric ecological interactions in aquatic ecosystems revealed through ultra-deep metagenomics. *Microbiome* 2019; **7**: 135.
6. Hyatt D, Chen G, LoCascio PF, Land ML, Larimer FW, Hauser LJ. Prodigal: prokaryotic gene recognition and translation initiation site identification. *BMC Bioinformatics* 2010; **11**.
7. Okazaki Y, Nishimura Y, Yoshida T, Ogata H, Nakano S. Genome-resolved viral and cellular metagenomes revealed potential key virus-host interactions in a deep freshwater lake. *Environmental Microbiology* 2019; **21**: 4740–4754.
8. Edwards RA, McNair K, Faust K, Raes J, Dutilh BE. Computational approaches to predict bacteriophage-host relationships. *FEMS Microbiology Reviews* 2016; **40**: 258–272.
9. Ghai R, Mehrshad M, Mizuno CM, Rodriguez-Valera F. Metagenomic recovery of phage genomes of uncultured freshwater actinobacteria. *The ISME Journal* 2017; **11**: 304–308.
10. Mann NH, Cook A, Millard A, Bailey S, Clokie M. Bacterial photosynthesis genes in a virus. *Nature* 2003; **424**: 741–741.
11. Rho M, Wu Y-W, Tang H, Doak TG, Ye Y. Diverse CRISPRs Evolving in Human Microbiomes. *PLOS Genetics* 2012; **8**: e1002441-.
12. Laslett D, Canback B. ARAGORN, a program to detect tRNA genes and tmRNA genes in nucleotide sequences. *Nucleic Acids Res* 2004; **32**: 11–16.
